# Direct RNA sequencing reveals selective remodeling of the host m6A epitranscriptome during *Leishmania* infection

**DOI:** 10.64898/2026.08.13.744690

**Authors:** Anelise Gonçalves Marino, Miguel Antonio do Nascimento Garcia, Bruno Souza Bonifácio, Carlos Eduardo Gomes Alves, Artur Honorato Reis, Paulo Otávio Lourenço Moreira, Marton Kaique de Andrade Cavalcante, Valéria Rêgo Alves Pereira, Maria Edileuza Felinto de Brito, Fabrício Castro Machado, Rafael de Freitas e Silva, Rubens Lima do Monte Neto, Elton J. R. Vasconcelos, Nilmar Silvio Moretti

## Abstract

N6-methyladenosine (m6A) is the most abundant modification in eukaryotic mRNA, yet its role in host responses to protozoan infection remains poorly explored. Here, we provide the first characterization of the host m6A epitranscriptome during *Leishmania amazonensis* infection, integrating clinical data, in vitro macrophage models, and direct RNA sequencing. Analysis of publicly available transcriptomic datasets from patients with cutaneous and visceral leishmaniasis, validated in an independent cohort of patients with cutaneous leishmaniasis, revealed coordinated remodeling of the m6A regulatory machinery, characterized by significant upregulation of the writer METTL3 and downregulation of the eraser ALKBH5 and the reader YTHDF3. These changes were recapitulated in RAW264.7 macrophages infected in vitro, where METTL3 protein levels increased progressively between 2 and 8 hours post-infection (hpi), while ALKBH5 abundance declined, resulting in elevated global m6A levels detectable as early as the first post-internalization time point. Transcriptome-wide mapping by direct RNA sequencing at 8 hpi identified 1,998 m6A-modified transcripts harboring 7,132 high-confidence sites in non-infected macrophages, compared with 2,956 modified transcripts and 13,495 sites in infected cells. Beyond this quantitative expansion, infection was associated with increased modification occupancy at shared sites, a greater prevalence of multi-site and multi-region methylation, and selective remodeling of immune-related and immunometabolic transcripts, including *Tlr4, Ccl2, Hmox1, Socs3, Rela, Hk1, Hk2, and Dicer1*. Altogether, these findings establish that *Leishmania* infection drives extensive yet selective remodeling of the host m6A landscape and position the host epitranscriptome as a previously unrecognized regulatory axis in *Leishmania*-macrophage interactions, with implications for the development of host-directed therapeutic strategies.

**Impact Statement:** N6-methyladenosine (m6A) is the most abundant modification in eukaryotic mRNA, and emerging evidence links epitranscriptomic remodeling to host responses against diverse pathogens. However, whether m6A regulates host-macrophage responses during Leishmania infection remains unknown. Using Oxford Nanopore Direct RNA sequencing, we provided the first transcriptome-wide map of host m6A modifications during *Leishmania amazonensis* infection. Infection selectively remodels the m6A landscape of macrophages, targeting immune-regulatory and immunometabolic transcripts, establishing the host epistranscriptome as a previously unrecognized regulatory layer in *Leishmania*-macrophage interactions.

## Introduction

Leishmaniases remain a major global health burden, affecting approximately 12 million people across 98 countries, with an estimated 1–1.5 million new cases of cutaneous leishmaniasis and 50,000–90,000 cases of visceral leishmaniasis reported annually (Pareyn *et al*., 2025). The diseases are caused by protozoan parasites of the genus *Leishmania*, which are transmitted to mammalian hosts through the bite of infected phlebotomine sandflies. Among the pathogenic species to humans, *Leishmania amazonensis* is a major etiological agent of cutaneous and diffuse cutaneous leishmaniasis in the Americas (Abadías-Granado *et al*., 2021). Infection by this parasite is frequently associated with chronic non-healing lesions characterized by high parasite loads and limited cell-mediated immune response, and impaired parasite clearance (Reithinger *et al*., 2007; Pareyn *et al*., 2025). These features make *L. amazonensis* a particularly relevant model for investigating host-parasite interactions and immune invasion mechanisms.

As obligate intracellular pathogen, *Leishmania* spp. parasites establish infection within macrophages, where they replicate inside specialized compartments known as the parasitophorous vacuoles. Macrophages therefore occupy a dual role during infection, serving both as a primary cellular niche that supports parasite survival and as key effector cells responsible for initiating and orchestrating antimicrobial immune response (Bogdan, 2020; Pareyn *et al*., 2025).

To establish and maintain a permissive intracellular niche, *Leishmania* parasites have evolved sophisticated strategies to manipulate host macrophage biology at multiple regulatory layers. These mechanisms include the suppression of reactive oxygen and nitrogen species production, modulation of cytokine signaling networks, impairment of antigen processing and presentation, and extensive reprogramming of host transcription responses. In particularly, *Leishmania* infection interferes with the activity of key transcription factors, including NF-κB, STAT1, IRF5, and HIF-1α, which are central regulators of macrophage activation, antimicrobial effector functions, and metabolic adaptations (Liu and Uzonna, 2012; Bogdan, 2020; Bichiou *et al*., 2021; Palomino-Cano *et al*., 2024).

Beyond transcriptional control, accumulating evidence indicates that *Leishmania* infection induces extensive post-transcriptional reprogramming of the host cell. In a seminal study, Chaparro et al. (Chaparro *et al*., 2020) showed that *L. donovani* infection triggers rapid and selective translational reprogramming in macrophages, altering the translation of distinct subsets of immune-related and metabolic mRNAs independently of changes in their transcript abundance. These findings highlight that the host cell proteome during *Leishmania* infection is shaped not only by transcriptional events but also by post-transcriptional and translational control mechanisms (Chaparro *et al*., 2020; Hagedorn *et al*., 2026). Despite these advances, the role of epitranscriptomic RNA modifications remains largely unexplored in the context of *Leishmania* infection, especially whether these modifications contribute to the dynamic regulation of host gene expression and macrophage response during parasite infection.

Epitranscriptomic modifications have emerged as a critical regulatory layer governing RNA metabolism and gene expression across eukaryotes (Boo and Kim, 2020; Cui *et al*., 2022). Among more than 170 known RNA modifications, N6-methyladenosine (m6A) is the most abundant internal modification in eukaryotic mRNA, present in approximately 25% of cellular transcripts (Zaccara *et al*., 2019). m6A is deposited co-transcriptionally at DRACH consensus motifs by a dynamic regulatory machinery comprising the methyltransferase “writer” complex (METTL3/METTL14, with accessory factors WTAP, VIRMA, ZC3H13, and RBM15), removed by demethylases “erasers” FTO and ALKBH5, while its biological effects are mediated by a diverse set of “reader” proteins such as YTHDF1-3, YTHDC1-2, and IGF2BP1-3 (Patil *et al*., 2018; Shi *et al*., 2019). Through the coordinated action of writers, erasers and readers, m6A regulates virtually all aspects of RNA metabolism, including transcript stability, splicing, nuclear export, and translation efficiency (Nachtergaele and He, 2018).

Critically, the functional consequences of m6A are highly context-dependent and determined by both the repertoire of reader proteins expressed and the positional distribution of methylation sites within transcripts. For example, YTHDF2 promotes mRNA decay through recruitment of the CCR4-NOT deadenylase complex, whereas YTHDF1 enhances cap-dependent translation, and YTHDF3 cooperates with both functions. In contrast, IGF2BP1-3 stabilize their target transcripts, meaning that the same m6A mark can produce opposing outcomes depending on which readers are expressed (Balacco and Soller, 2019; Shi *et al*., 2019; Zhou *et al*., 2024). Moreover, m6A sites located within 5′ UTR m6A can promote cap-independent translation under stress conditions, CDS methylation has been associated with translation-coupled mRNA degradation by YTHDF2, and 3′ UTR methylation serve as a major platform for reader-mediated regulation of transcript fate (Shi *et al*., 2019; Zhou *et al*., 2024).

The importance of m6A-mediated regulation and host-pathogen interactions has become increasingly evident in recent years. In macrophages, METTL3 deletion impairs host defense against *Salmonella typhimurium* in vivo, highlighting the essential role of m6A deposition in innate immune response (Tong *et al*., 2021). Conversely, the reader YTHDF2 acts as a negative regulator of LPS-induced inflammation by promoting the degradation of *Map2k4* and *Map4k4* mRNAs, thereby dampening MAPK signaling (Yu *et al*., 2019).

Also, accumulating evidence further indicates that diverse pathogens actively manipulate the host m6A machinery to create a cellular environment favorable for infection or to modulate the host defense. During SARS-CoV-2 infection, for example, the virus induces a global reduction of m6A in host transcripts while accumulating the modification on its own genome, suppressing cellular stress response and facilitating viral replication (Vaid *et al*., 2023). Similarly, infection of RAW264.7 macrophages with *Pseudomonas aeruginosa* promotes selective m6A deposition in 3′ UTRs of immune-related transcripts, including components of TLR4 pathway, with ALKBH5 acting as a key regulator of the innate immune response gene network (Feng *et al*., 2022). Evidence from protozoan infections also supports a pivotal role of epitranscriptomic regulation. In *Cryptosporidium parvum* infection, ALKBH5 downregulation via TLR/MyD88/NF-κB signaling drives global m6A elevation, promoting host epithelial defense (Xia *et al*., 2021a). Also, during *Toxoplasma gondii* infection of human macrophages, FTO-mediated demethylation of TNF-α mRNA, reduces YTHDF2-dependent transcript decay promoting host survival (Qin *et al*., 2025a). Altogether, these studies establish that epitranscriptomic remodeling is an important regulatory interface in host-pathogen interactions. However, whether similar mechanisms contribute to the host response against trypanosomatid parasites, particularly *Leishmania*, has not been addressed.

Recent advances in Oxford Nanopore direct RNA sequencing (DRS) have made possible to map RNA modifications at transcriptome-wide scale and at isoform resolution, without the antibody enrichment bias inherent to immunoprecipitation-based approaches such as m6A-seq and MeRIP-seq (Diensthuber and Novoa, 2025). Here, we apply this technology in combination with patient-derived transcriptomic data and controlled in vitro macrophage infection models to provide the first characterization of the host m6A epitranscriptome during *L. amazonensis* infection. Through an integrated analysis encompassing clinical cohorts, infection kinetics, and transcriptome-wide m6A mapping, we determine the landscape, selectivity, and immunological context of host epitranscriptomic remodeling during *Leishmania* infection and provide a high-resolution dataset as a valuable resource for future mechanistic investigation by the research community.

## Results

### Dysregulation of the m6A regulatory machinery in human leishmaniasis

To investigate whether components of the m6A regulatory machinery are altered during human leishmaniasis, we first analyzed three publicly available RNA-seq datasets generated from blood samples of patients with cutaneous or visceral leishmaniasis and skin lesions biopsies from individuals infected with *L. amazonensis* (Supplementary Table S1). Together, these datasets revealed infection-associated changes in the expression of multiple m6A regulatory factors, including writers, erasers and readers, although the magnitude and direction of these changes varied across clinical cohorts (Figure 1A-B and Supplementary Figure 1).

**Figure 1.**
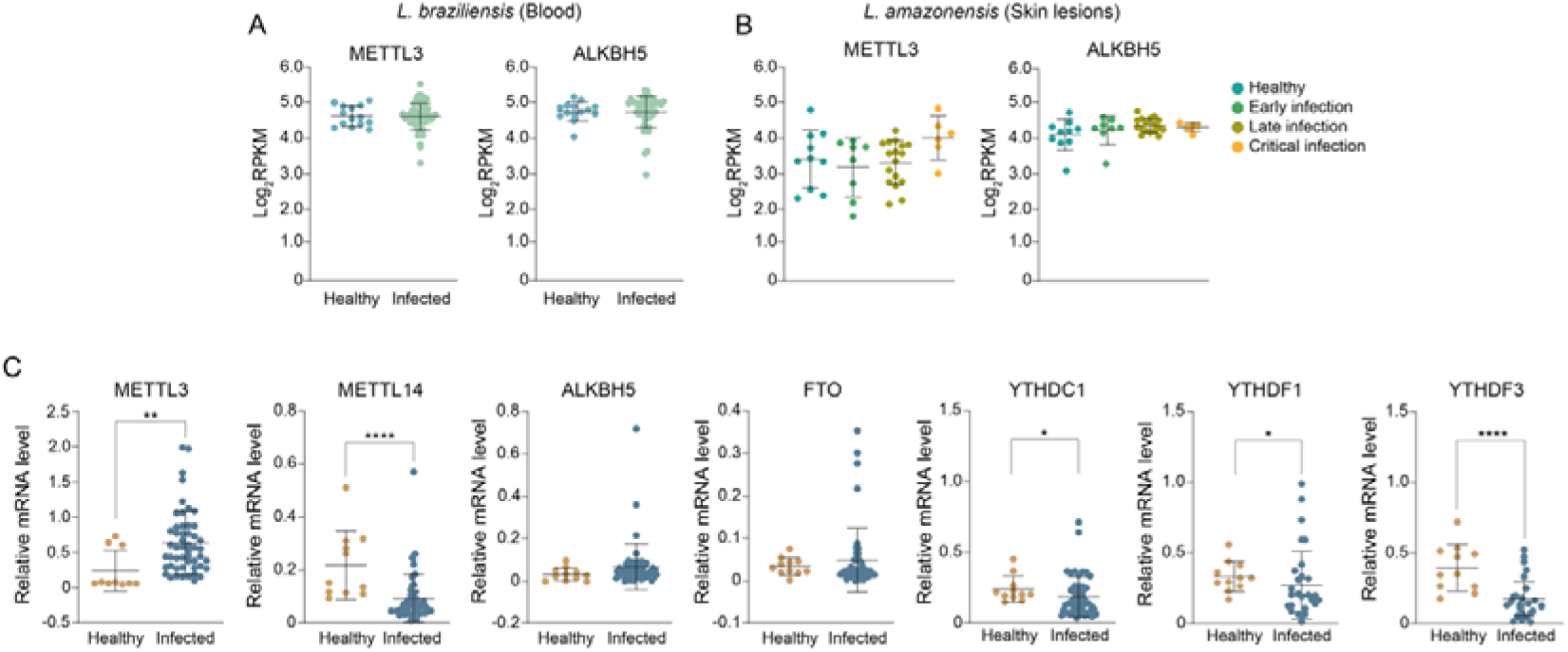
Remodeling of m6A regulatory machinery expression in human leishmaniases. **A.** Expression of the m6A writer METTL3 and the eraser ALKBH5 in whole-blood samples from healthy individuals and patients infected with *L. braziliensis*, based on publicly available RNA-seq data (GEO: GSE162760). **B.** Expression of METTL3 and ALKBH5 in skin lesion biopsies from patients infected with *L. amazonensis* stratified according to disease stage (early infection, late infection, and chronic infection), based on publicly available data (SRA BioProject: PRJNA307599). **C.** Relative mRNA expression of selected m6A regulatory factors (METTL3, METTL14, ALKBH5, FTO, YTHDC1, YTHDF1, and YTHDF3) in PBMCs from patients diagnosed with cutaneous leishmaniasis and healthy controls, determined by RT–qPCR. Gene expression levels were normalized to *ACTIN* and calculated using the ΔCt method. Data are presented as mean DSD. Statistical significance was assessed using the Mann–Whitney *U* test (\**p* < 0.05, \*\**p* < 0.01, \*\*\*\**p* < 0.0001).

Among the evaluated factors, the m6A writer METTL3 showed a tendency of increased expression in infected samples, particularly in *L. amazonensis* skin lesions, whereas the erasers ALKBH5 and FTO demonstrated reduced expression in specific datasets, although without reaching statistical significance (Figure 1A-B and Supplementary Figure 1). In addition, multiple m6A reader proteins exhibited differential expression patterns between cohorts, including the YTH domain-containing and IGF2BP families, supporting the notion that leishmaniasis is associated with broad remodeling of the host m6A regulatory machinery rather than uniform regulation of individual components.

To validate these observations in an independent clinical cohort, we quantified the expression of selected m6A regulators in PBMC samples obtained from 50 patients diagnosed with cutaneous leishmaniasis and 10 healthy controls. Consistent with the patterns observed in the transcriptomic datasets, METTL3 expression was significantly increased in infected individuals, confirming the association between infection and upregulation of m6A methylation machinery (Figure 1C). In contrast, the expression of METTL14, YTHDC1, YTHDF1 and YTHDF3 was significantly reduced in infected individuals, indicating concomitant alterations in both m6A deposition and recognition pathways. FTO expression remained largely unchanged between groups, whereas ALKBH5 displayed considerable inter-individual variability and did not reach the magnitude or consistency of differential expression observed for METTL3 (Figure 1C). Together, these results indicate that human leishmaniases is associated with extensive remodeling of the host m6A regulatory machinery, involving multiple components.

### METTL3 and ALKBH5 expression dynamics during macrophage infection

To determine whether the changes observed in patient-derived samples could be reproduced in a controlled experimental setting, RAW264.7 macrophages were infected with *L. amazonensis* stationary promastigotes, and the abundance of the m6A writer METTL3 and the eraser ALKBH5 was monitored over the course of infection.

Western blot analyses revealed modulation of METTL3 protein levels in infected macrophages, with increased abundance relative to non-infected controls observed from 0 to 4 hpi, while differences at 8 and 24 hpi were less pronounced, approaching baseline levels (Figure 2A). Immunofluorescence analysis, which enabled more robust quantification across a larger number of cells, revealed a more defined temporal pattern, with elevated METTL3 signal in infected cells at 2–8 hpi and reduced levels at 24 hpi (Figure 2B).

**Figure 2.**
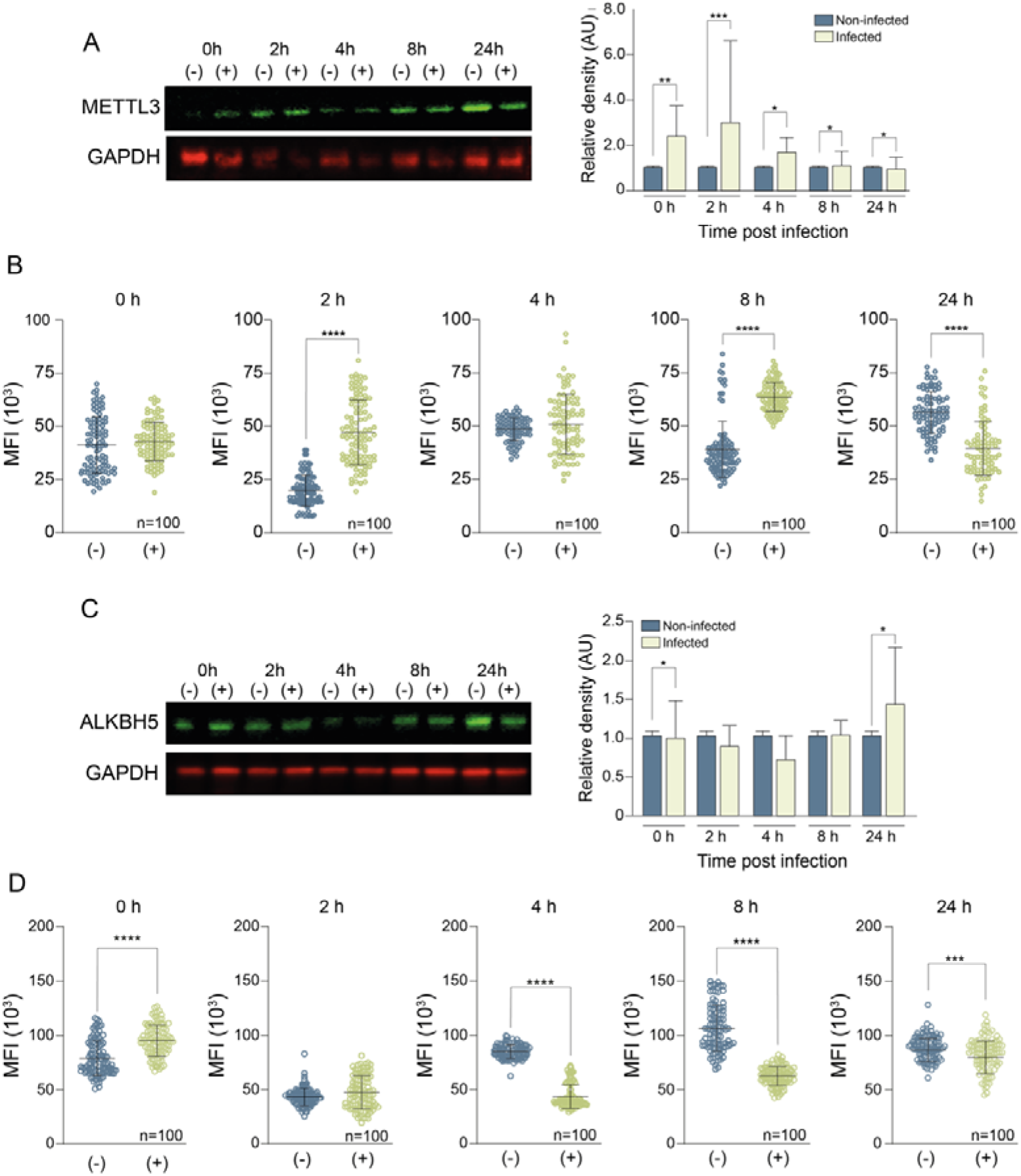
METTL3 and ALKBH5 protein expression during *L. amazonensis* infection in RAW264.7 macrophages. RAW264.7 macrophages were infected with stationary-phase *L. amazonensis* promastigotes, and protein levels were assessed at 0, 2, 4, 8, and 24 h post-infection. **A.** Representative western blot results of METTL3 protein levels and corresponding densitometric quantification. **B.** Quantification of METTL3 mean fluorescence intensity (MFI) determined by immunofluorescence analysis. **C.** Representative western blot showing ALKBH5 protein levels and corresponding densitometric quantification. **D.** Quantification of ALKBH5 mean fluorescence intensity (MFI) determined by immunofluorescence analysis. Confocal images were acquired at identical settings, and fluorescence was quantified at the whole-cell level in at least 100 cells per experimental condition. Statistical analysis was performed using a *t*-test (\*\*\**p* < 0.001, \*\*\*\**p* < 0.0001). The experiments were performed in triplicate (n = 3 independent experiments), and the images are representative of the results observed.

In contrast to METTL3, ALKBH5 displayed a distinct expression profile. Western blot analyses revealed a more heterogeneous pattern, with a significant increase in ALKBH5 abundance at 0 hpi, no significant differences at intermediate time points (2–8 hpi), and a significant increase again at 24 hpi (Figure 2C). Immunofluorescence analyses, further demonstrated dynamic changes in ALKBH5 abundance, characterized by a transient increase in signal intensity immediately following parasite-host contact (0 hpi), followed by progressive decline at subsequent time points. The reduction was particularly pronounced between 4 and 8 hpi, coinciding with the period of maximal METTL3 accumulation (Figure 2D).

Despite the infection-induced changes in protein abundance, confocal microscopy revealed no major changes in their intracellular distribution of either METTL3 or ALKBH5. Both proteins retained their predominant subcellular localization patterns through the infection course (Supplementary Figure 2 and 3).

**Figure 3.**
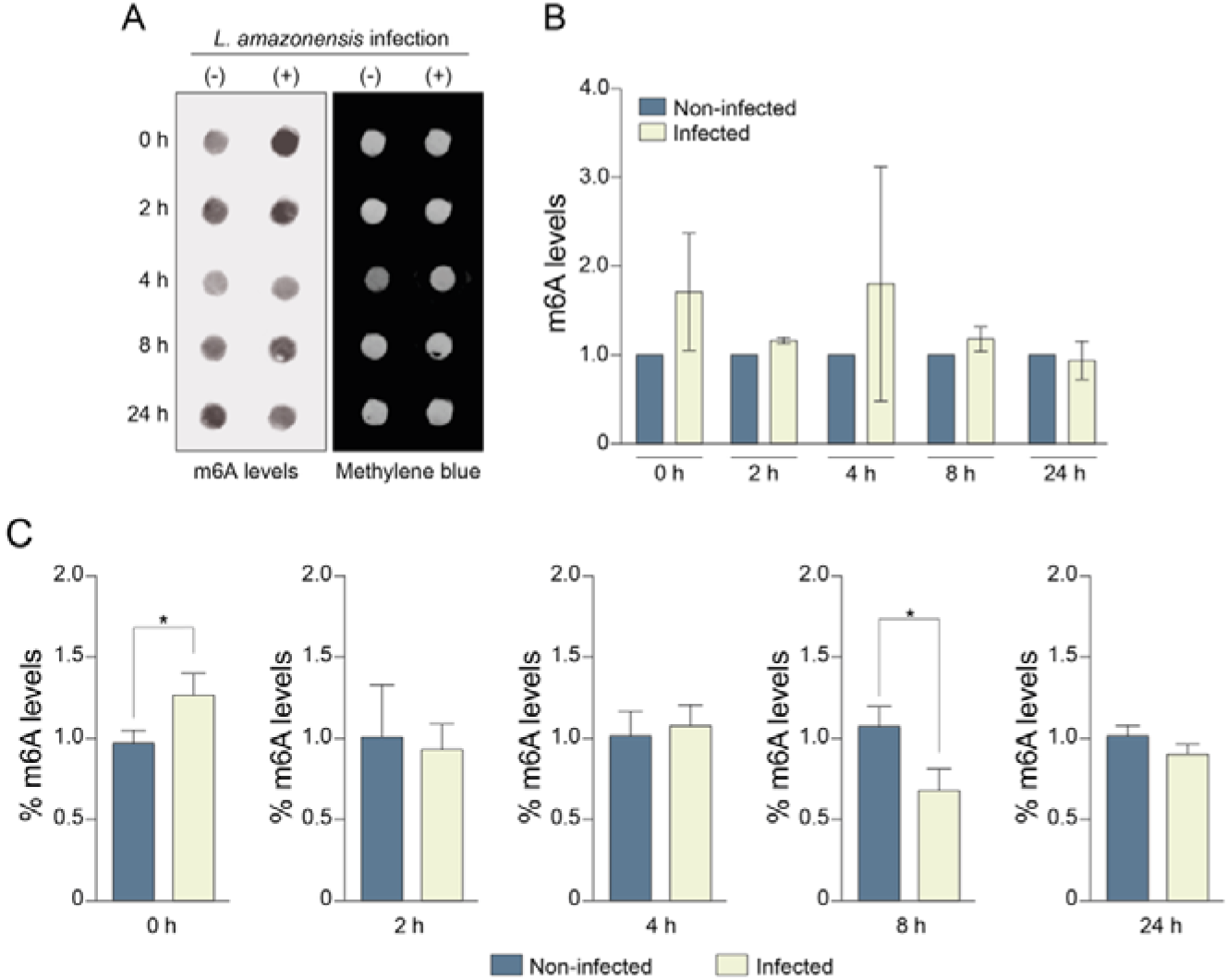
Changes in global m6A levels during *L. amazonensis* infection of macrophages. **A.** Dot blot analysis of total RNA isolated from RAW 264.7 macrophages infected with stationary-phase *L. amazonensis* promastigotes at 0, 2, 4, 8, and 24 hpi. Membranes were probed with anti-m6A antibody to measure global m6A levels, and methylene blue staining was used as a loading control. The 0 hpi time point corresponds to the completion of the initial 2 h parasite-macrophage cell interaction period and the removal of non-internalized parasites. **B.** Densitometric quantification of m6A signals obtained from dot blot analyses and normalized to the corresponding methylene blue staining. Values are presented relative to matched non-infected controls. **C.** Independent quantification of global m6A levels using the EpiQuik™ m6A RNA Methylation Quantification Kit (Epigentek) performed on the same RNA samples. Data are presented relative to the corresponding non-infected controls at each time point. Data are shown as mean SD from three independent experiments (n= 3). A representative dot blot image is shown in panel A. Statistical significance was determined using Student’s *t*-test (\**p* < 0.05).

We next examined whether the infection-associated changes observed at protein level were accompanied by alterations in transcript abundance. We quantified METTL3 and ALKBH5 mRNA levels during macrophage in vitro infection, and although both transcripts exhibited modest fluctuations, their expression profile did not fully mirror the corresponding protein dynamics (Supplementary Figure 4). It suggests that infection-associated changes in METTL3 and ALKBH5 abundance may involve additional regulatory mechanisms.

### *L. amazonensis* infection is associated with changes in global m6A levels

Given the infection-associated modulation of METTL3 and ALKBH5, we next sought to determine whether these changes translated into alterations in global m6A levels during macrophage infection. To address this question, total m6A abundance was quantified by Dot blot analyses at multiple time points followed *L. amazonensis* infection (Figure 3A). Notably, elevated m6A levels were already observed at the first post-infection time point (0 h), which corresponds to the completion of the initial 2 h parasite-host interaction period and the removal of non-internalized parasites. Increased m6A abundance was also detected at later early-stage time points, particularly at 4 hpi, followed by a gradual return to baseline levels by 24 hpi (Figure 3B). These findings demonstrated that *L. amazonensis* infection triggers an early and transient increase in host RNA methylation, consistent with the temporal modulation of key m6A regulatory factors observed in protein level.

To independently validate the dot blot results, global m6A levels were quantified using a colorimetric RNA methylation assay. Consistent with the dot blot data, infected macrophages exhibited elevated m6A levels at 0 hpi relative to non-infected controls (Figure 3C), confirming that infection rapidly induces changes in the host epitranscriptome following parasite internalization. In contrast, a significantly reduction in global m6A levels was observed at 8 hpi, whereas no significant differences were observed at 2, 4, or 24 hpi. Although the magnitude and timing of the changes varied between methodologies, both independent approaches consistently demonstrated that *L. amazonensis* infection is associated with dynamic remodeling of global m6A levels in host macrophages.

### Global profiling of m6A-modified transcripts during *L. amazonensis* infection by direct RNA sequencing

To investigate transcript-specific changes in m6A methylation during *L. amazonensis* infection, we performed direct RNA sequencing (DRS) on RAW264.7 macrophages collected at 8 hpi. This time point was selected based on the differences in global m6A levels observed between infected and non-infected cells (Figure 3). Three biological replicates were generated for each condition and analyzed using a Nanopore-based workflow optimized for transcriptome-wide m6A detection and quantification (Figure 4A).

**Figure 4.**
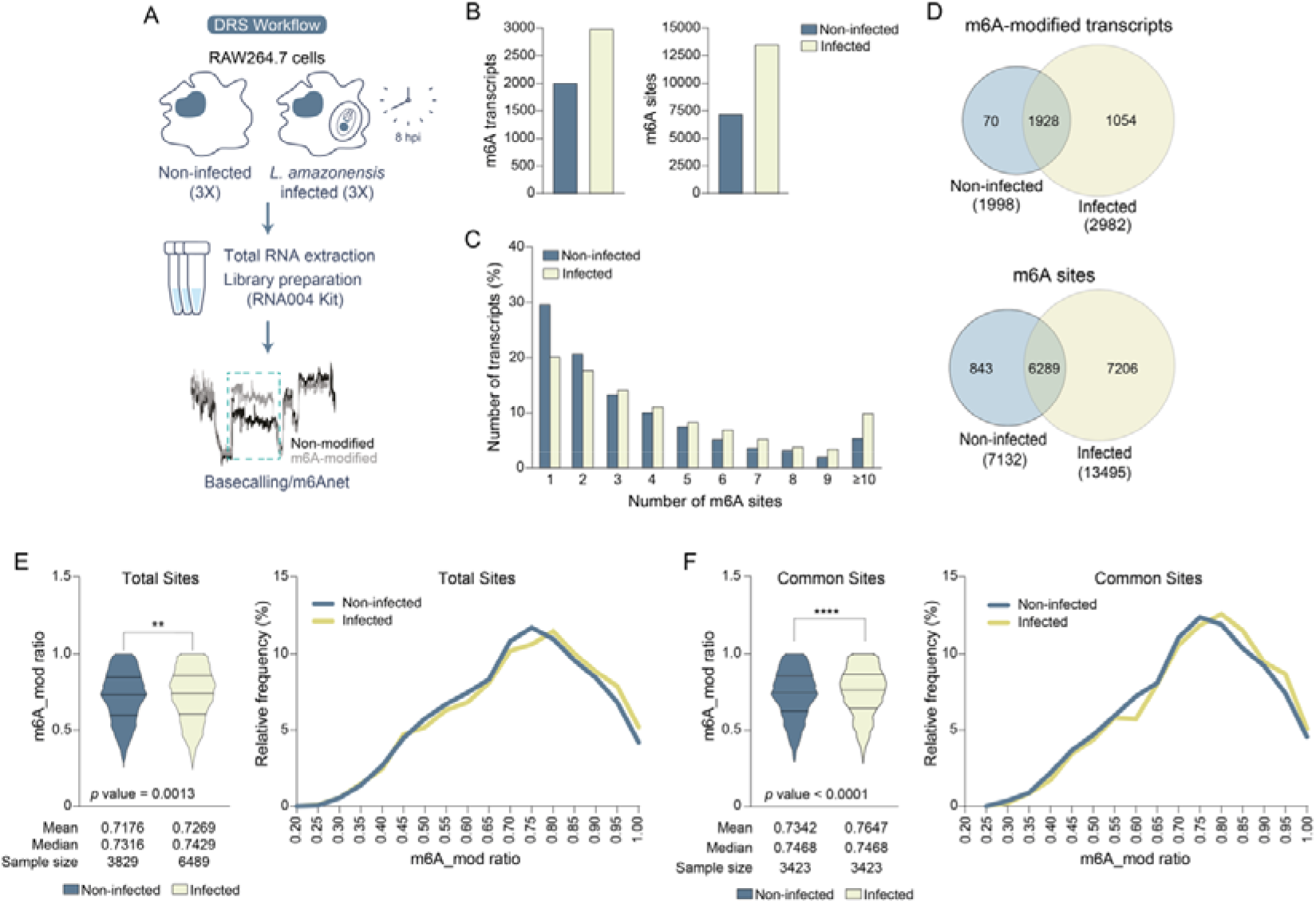
Transcriptome-wide profiling of m6A-modified transcripts during *L. amazonensis* infection by direct RNA sequencing. **A.** Schematic overview of the direct RNA sequencing (DRS) workflow used to identify m6A-modified transcripts in non-infected and *L. amazonensis*-infected RAW264.7 macrophages collected at 8 hpi. Three biological replicates were analyzed for each condition. Total RNA was subjected to direct RNA sequencing using the Oxford Nanopore platform, followed by m6A identification using the m6anet pipeline. **B.** Total number of m6A-modified transcripts (left) and high-confidence m6A sites (right) identified in non-infected and infected macrophages. **C.** Distribution of transcripts according to the number of detected m6A sites. Values represent the percentage of modified transcripts containing one or more m6A sites in each condition. **D.** Venn diagrams showing the overlap of m6A-modified transcripts (top) and m6A sites (bottom) between non-infected and infected macrophages. Shared and condition-specific transcripts and methylation sites are indicated. **E.** Global distribution of m6A mod ratio considering all detected m6A sites after filtering the sites for their presence in at least two of the three biological replicates per condition. Violin plots summarize the distribution of occupancy values, and frequency histograms show the relative abundance of sites across different m6A mod ratio intervals. Statistical differences between distributions were assessed using the Kolmogorov–Smirnov test (*p* = 0.0013). **F.** Distribution of m6A modification ratios considering only m6A sites shared between non-infected and infected macrophages. Violin plots and frequency histograms are shown as in panel E. Statistical differences were assessed using the Wilcoxon matched-pairs signed-rank test (\*\*\*\**p* < 0.0001).

Following stringent quality filtering, high-confidence m6A site identification and integration of replicate datasets, we identified 1,998 m6A-modified transcripts containing a total of 7,132 m6A sites in non-infected macrophages. In contrast, infected-macrophages exhibited 2,982 methylated transcripts harboring 13,495 m6A sites (Figure 4B and Supplementary Tables S3-S5).

We next examined the distribution of m6A sites across individual transcripts. Consistent with previous transcriptome-wide m6A studies most modified transcripts in both conditions contained a single m6A site (Figure 4C) (Dominissini *et al*., 2012; Meyer *et al*., 2012). However, infected macrophages exhibited an apparent shift toward greater methylation complexity, characterized by an increase proportion of transcripts harboring multiple m6A sites, particularly those containing ten or more modifications. This pattern suggests that *L. amazonensis* infection not only expands the repertoire of methylated transcripts but also enhances the degree of methylation within individual transcripts.

To assess the extent of overlap between conditions, we compared both m6A-modified transcripts and individual m6A sites between non-infected and infected macrophages. At the transcript level, most modified transcripts were shared between conditions (1,928 transcripts), indicating that much of the basal m6A landscape is maintained during infection. However, infection was associated with the appearance of 1,054 uniquely methylated transcripts, whereas only 70 modified transcripts were unique to non-infected controls (Figure 4D). Analysis at the single-site resolution revealed even a more extensive remodeling. Although 6,289 m6A sites were shared between both conditions, infected macrophages contained 7,206 unique m6A sites, compared with only 843 sites detected exclusively in non-infected cells (Figure 4D). These findings indicate that *L. amazonensis* infection is associated with both the emergence of newly methylated transcripts and extensive reorganization of methylation sites within transcripts that remain methylated under both conditions.

We next evaluated the degree of methylation at individual sites by analyzing the m6A modification ratio (m6A_mod ratio), a metric that estimates the proportion of sequencing reads carrying an m6A modification at a given nucleotide position. To ensure robust occupancy estimates, this analysis was restricted to high confidence m6A sites detected in at least two of the three biological replicates per condition. Analysis of all detected m6A sites revealed an increase modification occupancy in infected macrophages (Mean: 0.7269) compared with non-infected controls (Mean: 0.7176) (Figure 4E). This global shift was accompanied by an increase frequency of highly occupied sites in infected cells, as shown by the distribution of m6A modification ratio values (Figure 4E, right panel).

To determine whether this effect was restricted to newly acquired sites or also affected pre-existing methylation events, we focused the analysis on m6A positions shared between infected and non-infected macrophages. Even under this more stringent criterion, infected cells displayed higher m6A occupancy than controls (Mean: 0.7647 in infected versus Mean: 0.7342 in non-infected cells) (Figure 4F), indicating that infection is associated not only with the emergence of additional m6A sites but also with increased methylation occupancy at sites that remain modified under both conditions.

Collectively, these results demonstrate extensive transcriptome-wide remodeling of host m6A methylation during *L. amazonensis* infection, characterized by an expanded repertoire of methylated transcripts, the emergence of numerous infection-specific m6A sites, and elevated modification occupancy.

### *L. amazonensis* infection increases multi-region m6A modification of host transcripts

Because the biological consequences of m6A is strongly influenced by its positional context within transcripts, we next investigated the regional distribution of m6A sites across transcript architecture during *L. amazonensis* infection. Specifically, we quantified the number of m6A-modified transcripts containing at least one methylation site within the 5′ UTR, the CDS, and the 3′ UTR. In both non-infected and infected conditions most of m6A deposition was detected within the CDS and 3′ UTR regions, whereas relatively few transcripts had m6A sites within 5’ UTR (Figure 5A). Infection increased the number of modified transcripts over all transcript regions, with the most pronounced expansion observed in the CDS and 3’ UTR regions (Figure 5A).

**Figure 5.**
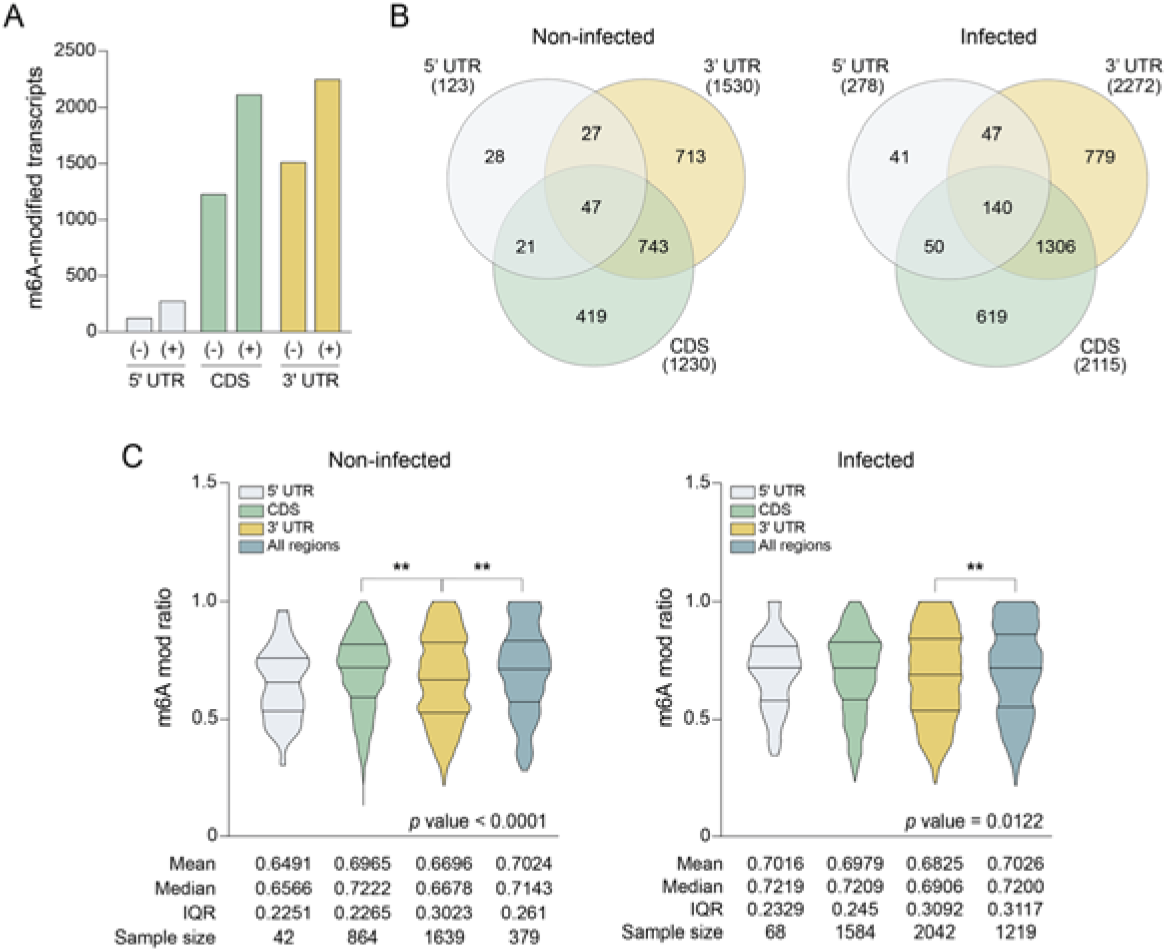
Regional distribution of m6A modifications across transcript regions in non-infected and *L. amazonensis*-infected macrophages. **A.** Number of m6A-modified transcripts containing at least one m6A site within 5’ UTR, CDS and 3’ UTR in non-infected (-) and infected (+) macrophages. **B.** Venn diagrams showing the overlap of m6A-modified transcripts among the 5′ UTR, CDS, and 3′ UTR regions in non-infected (left) and infected (right) macrophages. Numbers indicate transcripts exclusively modified within a single region or shared among multiple transcript regions. **C.** Distribution of m6A modification ratios (m6A mod ratio) for m6A sites located within the 5′ UTR, CDS, 3′ UTR, or in transcripts harboring modifications in all three regions. Violin plots are shown separately for non-infected (left) and infected (right) macrophages. Mean, median, interquartile range (IQR), and sample size are indicated below each plot. Statistical analyses were performed using the Kruskal–Wallis test followed by Dunn’s multiple-comparison test. Significant differences are indicated by asterisks (**p < 0.01).

Also, we examined the overlap of m6A-modified transcripts among distinct transcript regions. In non-infected macrophages, 47 transcripts harbored m6A sites simultaneously within the 5′ UTR, CDS, and 3′ UTR (Figure 5B). Following infection, this number increased nearly threefold to 140 transcripts (Figure 5B). Similar scenario was observed for transcripts harboring m6A sites in both CDS and 3’ UTR, whose number increased from 743 in non-infected cells to 1,306 in infected cells. These findings indicate that *L. amazonensis* infection not only increases the number of methylated transcripts but also enhances the architectural complexity of m6A deposition, resulting in a greater proportion of transcripts bearing modifications across multiple functional regions.

We next compared the m6A occupancy using the m6A mod ratio values among distinct transcript regions. In non-infected macrophages, m6A sites located within the CDS (n = 864) and transcripts harboring m6A modifications across all three regions (n = 379) showed highest median m6A mod ratio values (0.7222 and 0.7143, respectively) whereas the 5’ UTR showed comparatively lower occupancy levels (Median: 0.6566) (Figure 5C and Supplementary Figure 5). In addition, the 3’ UTR displayed the greatest variability in m6A occupancy, as indicated by its larger interquartile ranges (IQR = 0.3023), suggesting increased heterogeneity of modification levels within this region (Figure 5C and Supplementary Figure 5).

A broadly similar pattern was observed in infected macrophages. M6A sites located within CDS (n = 1584), 3′ UTR (n = 2042), and multi-region-associated transcripts (n = 1219) exhibited comparable median m6A occupancy values around 0.72 (Figure 5C). However, dispersion analysis revealed important differences between these categories. While the CDS maintained a relatively narrow interquartile range (IQR = 0.245), indicating a more stable distribution of m6A levels, both the 3′ UTR (IQR = 0.3092) and multi-region-associated transcripts (IQR = 0.3117) showed increased variability, consistent with a more heterogeneous distribution of m6A mod ratio values (Figure 5C and Supplementary Figure 5).

To further characterize the spatial organization of m6A deposition, we analyzed the positional distribution of m6A sites relative to translation initiation and termination codons. Although the global positional profiles remained broadly similar between the conditions, infected macrophages showed a modest increase in the density of m6A sites near the translation initiation codon (Supplementary Figure 5C), whereas accumulation near the stop codon was less pronounced compared to non-infected cells (Supplementary Figure 5D).

### Selective remodeling of m6A occupancy at conserved transcript sites during infection

After characterizing the regional distribution of m6A-modified transcripts, we next investigated whether *L. amazonensis* infection altered m6A occupancy at individual methylation sites. To this end, we focused on m6A sites shared between non-infected and infected macrophages within each transcript region.

Consistent with the global expansion of the methylome observed during infection, the total number of detected m6A sites increased substantially in all transcript regions, including the 5′ UTR, CDS, and 3′ UTR (Figure 6A). Despite this increase, a large proportion of m6A sites remained conserved between conditions, with 160, 2,590, and 3,539 shared sites identified in the 5′ UTR, CDS, and 3′ UTR, respectively (Figure 6B).

**Figure 6.**
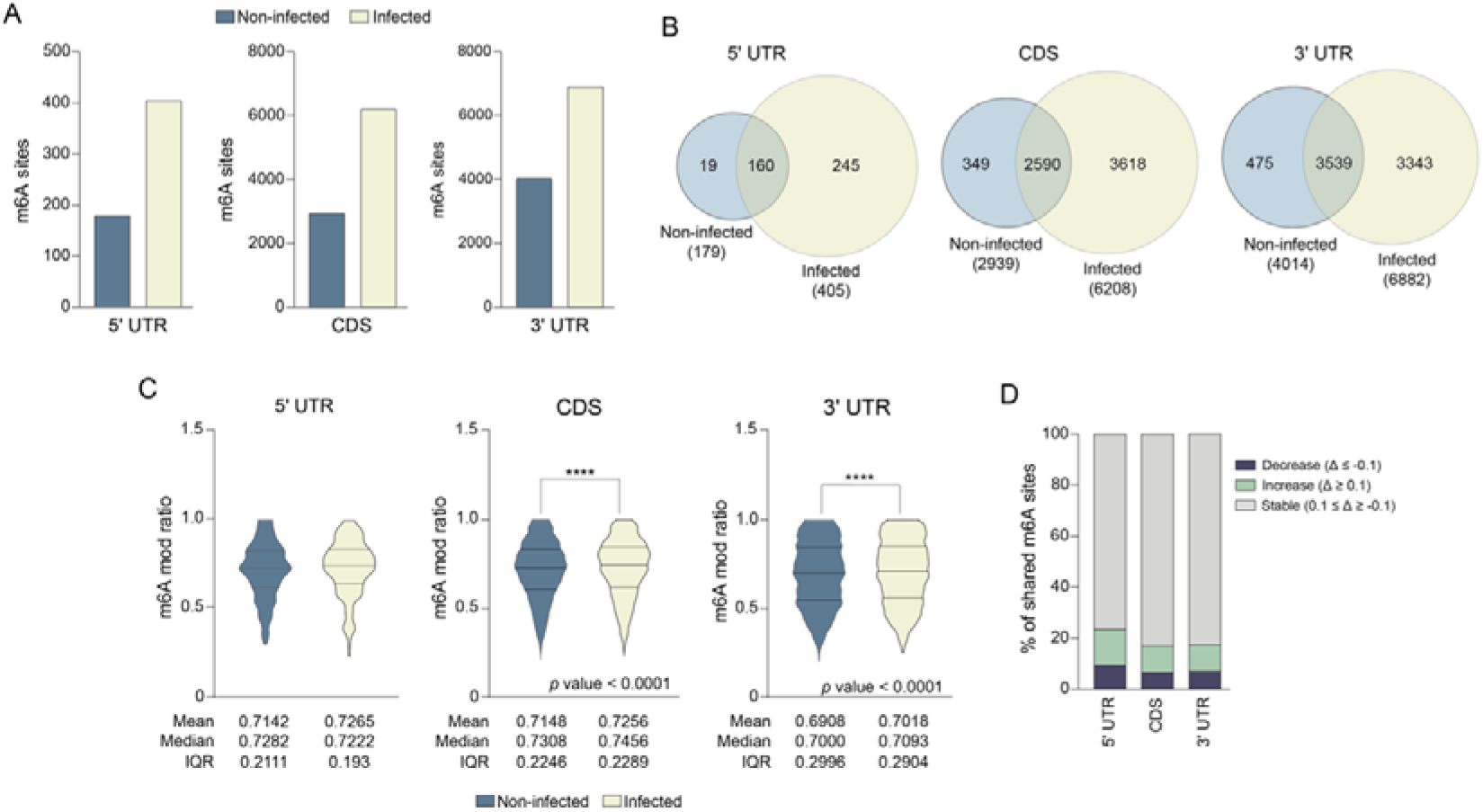
Selective remodeling of m6A occupancy at conserved transcript sites during *L. amazonensis* infection. **A.** Number of m6A sites detected within the 5′ UTR, CDS, and 3′ UTR of transcripts from non-infected and *L. amazonensis*-infected macrophages. **B.** Venn diagrams showing the overlap of m6A sites identified in non-infected and infected macrophages within each transcript region (5′ UTR, CDS, and 3′ UTR). Numbers indicate region-specific and shared m6A sites between conditions. **C.** Distribution of m6A mod ratio for m6A sites shared between non-infected and infected macrophages within the 5 ′UTR, CDS, and 3′ UTR. Violin plots display the distribution of m6A mod ratio values, with mean, median, and interquartile range (IQR) indicated below each plot. Statistical comparisons were performed using paired Wilcoxon matched-pairs tests. Significant differences were observed for CDS and 3′ UTR sites (\*\*\*\**p* < 0.0001), whereas no significant difference was detected for 5′ UTR sites. D. Classification of shared m6A sites according to changes in m6A occupancy between infected and non-infected macrophages (Δm6A mod ratio = infected − non-infected). Sites were categorized as increased (Δm6A mod ratio ≥ 0.1), stable (−0.1 < Δm6A mod ratio < 0.1), or decreased (Δm6A mod ratio ≤ −0.1). Across all transcript regions, most shared sites remained stable, while a subset exhibited increased or decreased m6A occupancy during infection.

To determine whether infection influenced the degree of methylation at these conserved positions, we compared m6A mod ratio between infected and non-infected macrophages for shared sites within each transcript region. Sites located within the CDS and 3′ UTR showed higher m6A occupancy in infected macrophages (CDS mean: 0.7256; 3’ UTR mean: 0.7018) compared with uninfected controls (CDS median: 0.7148; 3’ UTR median: 0.6908) (Figure 6C). In contrast, no significant difference was observed for shared 5′ UTR sites (Figure 6C). Despite reaching statistical significance, the overall occupancy profiles were largely similar between infected and non-infected macrophages, with only subtle shifts across m6A modification ratio intervals (Supplementary Figure 5E).

To further quantify infection-associated epitranscriptomic remodeling, we classified shared m6A sites according to changes in modification occupancy, calculated as the difference in m6A modification ratio between infected and non-infected macrophages (Δm6A mod ratio = infected – non-infected). Across all transcript regions, most of shared sites exhibited relatively stable methylation levels (−0.1 < Δm6A mod ratio < 0.1), accounting for approximately 76-83% of all conserved m6A sites (Figure 6D). Nevertheless, a consistent subset of m6A conserved sites exhibited increased methylation occupancy during infection (Δm6A mod ratio ≥ 0.1), whereas only a small fraction showed reduced occupancy (Δm6A mod ratio ≤ -0.1) (Figure 6D).

Together, these findings indicate that *L. amazonensis* infection promotes selective remodeling of the host transcripts m6A occupancy at conserved methylation sites, particularly within the CDS and 3′ UTR.

### Functional annotation of infection-associated m6A-modified transcripts

To gain insight into the biological processes associated with m6A-modified transcripts, we performed functional enrichment analyses using genes corresponding to all m6A-modified transcripts identified in non-infected and infected macrophages (Supplementary Table S4 and S5). In both conditions, the most significantly enriched KEGG pathways were predominantly associated with RNA metabolism and post-transcriptional gene regulation, including spliceosome, RNA transport, and protein processing in the endoplasmic reticulum (Supplementary Figure 6A and B), consistent with the established roles of m6A in RNA homeostasis.

Although many pathways were shared between conditions, *L. amazonensis* infection was associated with an expanded representation of pathways related to intracellular trafficking, cellular stress adaptation, and host response to infection, such as endocytosis and mRNA surveillance (Supplementary Figure 6B). These findings indicate that infection-associated m6A remodeling extends beyond constitutive RNA-processing functions and encompasses transcripts involved in diverse cellular processes.

To further characterize the increased complexity of the host m6A epitranscriptome during infection, we identified transcripts harboring the greatest number of modification sites. Several transcripts exhibited substantially increased numbers of m6A sites in infected macrophages, including Ahnak, Dicer1, Tlr13, Ranbp2, Nup153, and Mki67 (Supplementary Figure 6C). Among them, Dicer1 has no detectable m6A sites in non-infected macrophages but emerged as one of the most highly modified transcripts in the infection condition, illustrating extensive remodeling of methylation patterns at specific transcripts.

Moreover, transcripts were classified according to m6A modification occupancy as stoichiometrically hypomethylated (m6A mod ratio ≤0.35) or hypermethylated (m6A mod ratio ≥ 0.75). While hypomethylated sites presented similar occupancy distributions among the conditions, infected macrophages have a larger number of hypermethylated sites (Supplementary Figure 6D), consistent with the global increase in m6A occupancy observed in Figure 4.

Finally, considering the significant increase in the number of transcripts bearing m6A simultaneously within the 5′ UTR, CDS, and 3′ UTR regions in infected macrophages, we sought to determine whether this subset of extensively modified transcripts was associated with specific biological functions. To this end, we performed Gene Ontology (GO) enrichment analysis using transcripts containing m6A sites across all three transcript regions. In non-infected macrophages, these transcripts were primarily enriched for processes related to cellular homeostasis, cytoskeletal organization, responses to ions, and regulation of protein modification (Supplementary Figure 6E), consistent with their involvement in maintaining basal cellular functions. In contrast, the corresponding transcripts set from infected macrophages exhibited a distinct functional profile, showing preferentially enrichment for pathways associated with metabolic regulation, macromolecule biosynthesis, cellular adaptation, and myeloid leukocyte differentiation (Supplementary Figure 6F).

### Immune-related transcripts undergo m6A remodeling during *L. amazonensis* infection

To investigate whether infection-associated m6A remodeling preferentially targets genes involved in immunity, we generated a curated panel of immune-related genes implicated in the macrophage response to *Leishmania* infection through a systematic review of the literature. This curated dataset comprised 333 genes corresponding to 1,483 annotated transcripts (Figure 7A and Supplementary Table S6), encompassing key regulators of innate immune signaling, inflammatory responses, antigen presentation, antimicrobial defense and macrophage activation. We then interrogated our DRS dataset to identify m6A-modified transcripts within this immune-related gene set.

**Figure 7.**
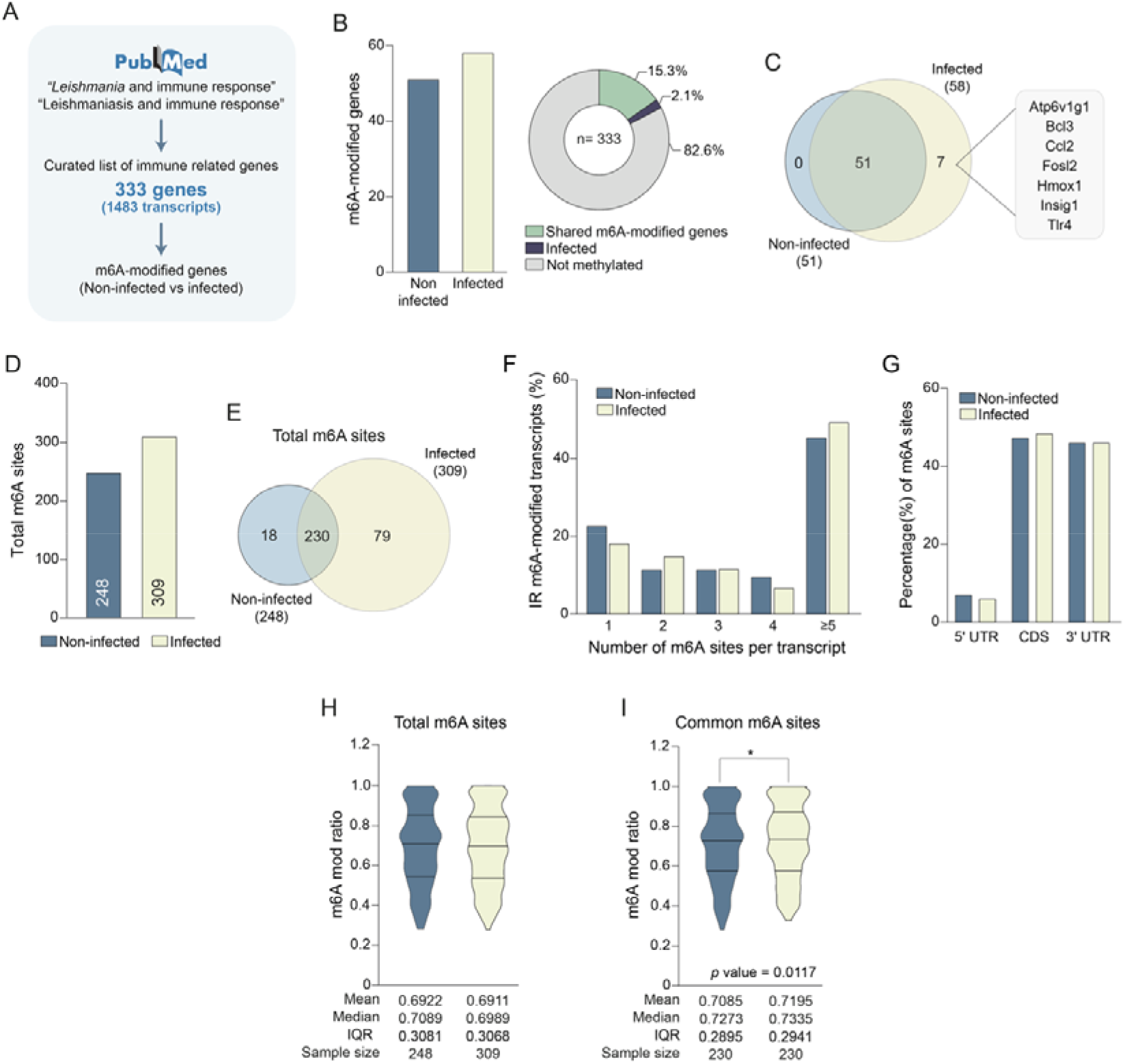
Immune-related transcripts exhibit infection-associated m6A remodeling during *L. amazonensis* infection. **A.** Workflow used to identify immune-related genes associated with *Leishmania* infection through literature curation, resulting in a dataset of 333 genes represented by 1,483 transcripts. **B.** Number and proportion of immune-related genes associated with m6A-modified transcripts in non-infected and infected macrophages. The donut chart summarizes the percentage of genes carrying shared or infection-specific m6A modifications within the curated dataset. **C.** Venn diagram showing the overlap of m6A-modified immune-related genes between non-infected and infected macrophages. The seven genes exclusively associated with m6A modifications in infected macrophages are indicated. **D.** Total number of m6A sites identified within immune-related transcripts in non-infected and infected macrophages. **E.** Venn diagram showing the overlap of m6A sites detected in immune-related transcripts between conditions. Numbers indicate shared and condition-specific m6A sites. **F.** Distribution of immune-related m6A-modified transcripts according to the number of m6A sites per transcript. Values are expressed as the percentage of modified transcripts within each category. **G.** Regional distribution of m6A sites across transcript regions (5′ UTR, CDS, and 3′ UTR) in immune-related transcripts from non-infected and infected macrophages. **H.** Distribution of m6A mod ratio considering all m6A sites identified in immune-related transcripts from non-infected and infected macrophages. Mean, median, interquartile range (IQR), and sample size are indicated below the violin plots. **I.** Comparison of m6A mod ratio restricted to m6A sites shared between non-infected and infected macrophages. Statistical significance was assessed using a paired Wilcoxon matched-pairs test (*p* = 0.0117).

Among these curated immune-related genes, 51 were associated with m6A-modified transcripts in non-infected samples, whereas 58 genes were modified in infected cells (Figure 7B). Most of these genes were shared between the conditions, with 51 common m6A-modified immune genes, corresponding to 15.3% of the curated dataset. In contrast, only seven genes (2.1%) were exclusively modified following infection (Figure 7B and C). The infection-specific m6A-modified genes comprised Atp6v1g1, Bcl3, Ccl2, Fosl2, Hmox1, Insig1, and Tlr4, all of which have previously been implicated in inflammatory signaling, stress responses, metabolic regulation, or innate immune activation (Figure 7C).

We next examined the organization of m6A sites within immune-related transcripts. A total of 248 high-confident m6A sites were detected within immune-associated transcripts in non-infected macrophages, whereas 309 sites were detected in infected ones (Figure 7D). Although most sites were shared between conditions (n=230), infected macrophages showed a larger number of condition-specific sites (n=79) than non-infected cells (n=18) (Figure 7E), suggesting that infection promotes the emergence of new methylation events within immune associated transcripts.

To further investigate the methylation complexity, we analyzed the number of m6A sites per transcripts. In both conditions, a large proportion of transcripts harbored five or more m6A-sites, although infected macrophages exhibited a modest increase in transcripts harboring multiple modification sites, accompanied by a reduction in transcripts containing only a single m6A site (Figure 7F). These findings suggest that infection-associated remodeling preferentially affects transcripts already enriched for m6A modifications.

Analysis of regional distribution of m6A sites revealed a broadly conserved pattern between conditions, with most sites localized within the CDS and 3’ UTR regions, and relatively few sites detected within the 5’ UTR (Figure 7G). This indicates that the infection-associated remodeling observed in immune-related transcripts occurs largely within the canonical transcript regions typically enriched for m6A deposition.

Finally, we evaluated whether *L. amazonensis* infection altered m6A occupancy by comparing m6A mod ratio values between conditions. Considering all detected sites, no significant differences in global occupancy were observed between non-infected and infected macrophages (Figure 7H). However, when the analysis was restricted to sites shared between conditions, infected macrophages displayed a modest but significant increase in m6A occupancy relative to non-infected controls (Wilcoxon signed-rank test, *p* = 0.0117; Figure 7I). These findings indicate that infection not only alters the repertoire of modified sites but also subtly increases methylation levels at conserved positions (Supplementary Table S7).

### m6A architecture changes of shared immune transcripts during infection

Although most m6A-modified immune-related genes were shared between non-infected and infected macrophages (Figure 7), it remained unclear whether *L. amazonensis* infection altered the internal organization of m6A sites within these transcripts. To address this question, we restricted the analysis to on immune-related transcripts that were m6A-modified under both conditions and examined how infection affected the site number, positional distribution and occupancy of individual m6A sites. Among these transcripts, 230 m6A sites were conserved in both conditions, whereas 18 and 64 sites were detected in non-infected and infected macrophages, respectively (Figure 8A). More than half of shared transcripts (56%) gained additional m6A sites during infection; 30% remained unchanged and only 13% exhibited site loss (Figure 8B).

**Figure 8.**
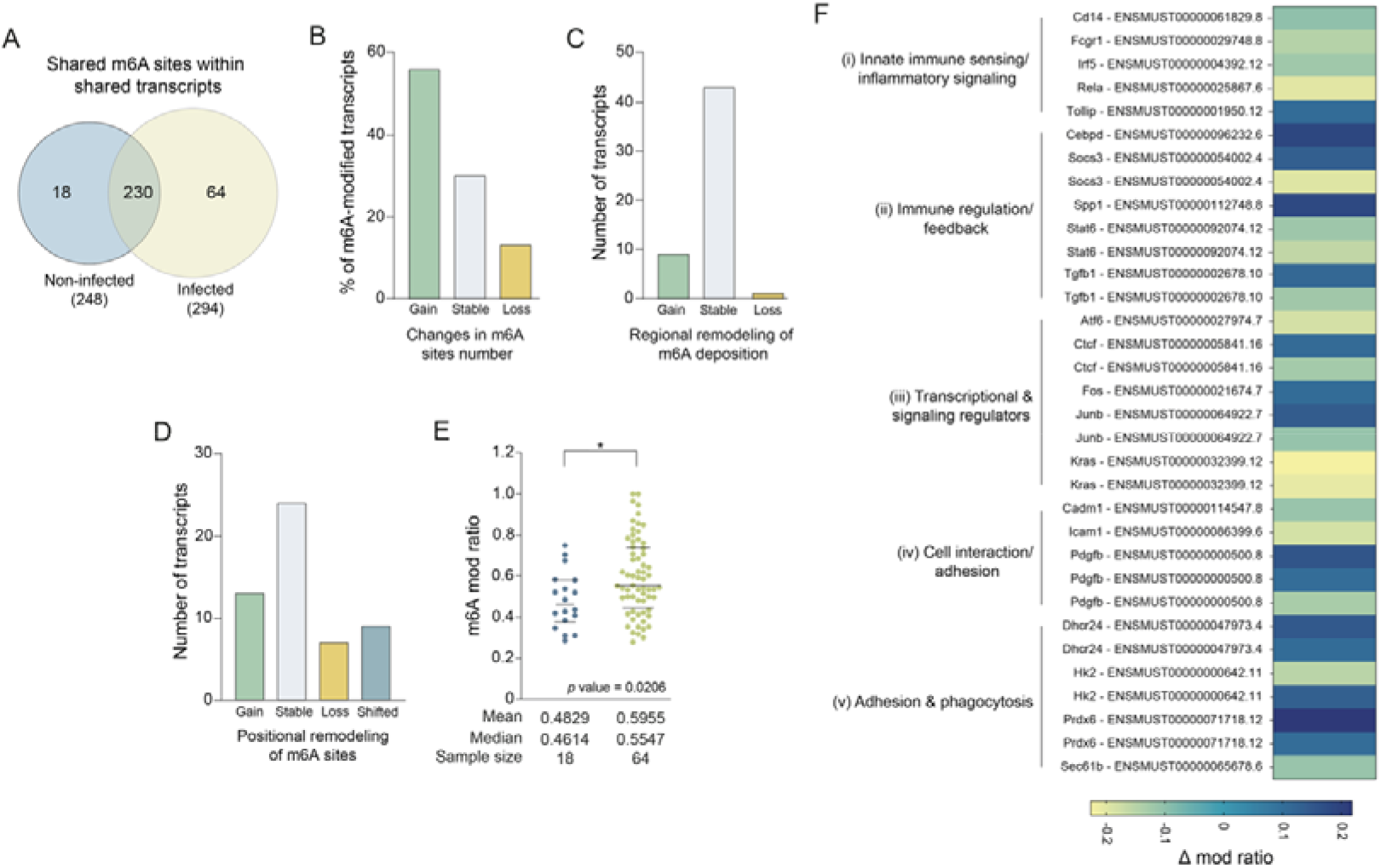
Site-specific alterations of m6A architecture within shared immune-related transcripts during infection. **A.** Venn diagram showing shared and condition-specific m6A sites identified within immune-related transcripts modified in both non-infected and infected macrophages. **B.** Distribution of shared immune-related transcripts according to changes in the number of m6A sites during infection. Transcripts were classified as exhibiting site gain, site loss, or no change (stable). **C.** Remodeling of transcript-region occupancy. Shared transcripts were classified according to changes in the transcript regions harboring m6A modifications (5′ UTR, CDS, and/or 3′ UTR), including gain of modification in previously unmodified regions, loss of regional occupancy, or no change (stable). **D.** Positional remodeling of m6A sites within shared transcripts. Transcripts were classified as exhibiting site gain, site loss, positional shifts (simultaneous gain and loss of sites), or no detectable changes (stable). **E.** Comparison of m6A occupancy (m6A mod ratio) between condition-specific sites detected exclusively in non-infected or infected macrophages. Horizontal lines indicate mean values. Statistical significance was assessed using the Mann–Whitney test (*p* = 0.0206). **F.** Heatmap showing transcripts with pronounced changes in Δm6A mod ratio among shared m6A sites. Transcripts were grouped into functional categories associated with host immune responses, including (i) innate immune sensing and inflammatory signaling, (ii) immune regulation and feedback mechanisms, (iii) transcriptional and signaling regulators, (iv) cell interaction and adhesion, and (v) adhesion and phagocytosis. Color scale indicates the magnitude and direction of Δm6A mod ratio values.

To determine if these additional sites altered the regional organization of m6A deposition we examined changes in transcript-region occupancy considering the 5’ UTR, CDS and 3’ UTR. Most transcripts (43 of 53) maintained the same combination of modified transcripts regions between conditions, whereas nine transcripts gained m6A deposition in previously unmodified regions and a single transcript exhibited regional loss (Figure 8C). Thus, despite the increase in m6A-site number, the overall regional topology of methylation remained largely stable during infection.

In contrast, analysis at single-site resolution revealed extensive remodeling of m6A deposition in immune-related transcripts. Only 24% of transcripts retained an identical pattern of m6A positions between conditions, whereas the majority showed evidence of remodeling, including acquisition of novel sites (n=13), loss of existing sites (n=7), or simultaneous gain and loss events resulting in positional shifts of m6A deposition (n=9) (Figure 8D). These findings indicate that infection predominantly reshapes the local architecture of m6A deposition without substantially altering the transcript regions targeted by methylation.

We next assessed if condition-specific sites differed in their degree of methylation. Sites detected exclusively in infected macrophages showed significantly higher m6A occupancy than sites identified only in non-infected cells (mean: 0.5955 versus 0.4829, respectively) (Figure 8E). This observation suggests that newly acquired sites are not merely stochastic methylation events but tend to exhibit relatively robust occupancy.

To explore the biological relevance of infection-associated m6A remodeling, we calculated the Δm6A values (infected – non-infected) for shared sites and focused on transcripts with the more pronounced occupancy variations (Δ ≥0.1 or ≤ -0.1). Functional classification revealed genes involved in multiple aspects of host defense, including innate immune sensing and inflammatory signaling (*Cd14*, *Fgr1*, *Irf5*, *Rela*, and *Tollip*), immune regulation and feedback mechanisms (*Cebpd*, *Socs3*, *Spp1*, *Stat6*, and *Tgfb1*), transcriptional and signaling regulation (*Atf6*, *Ctcf*, *Fos*, *Junb*, and *Kras*), cell interaction and adhesion pathways (*Cadm1*, *Icam1*, and *Pdgfb*), and adhesion/phagocytosis-associated functions (*Dhcr24*, *Prdx6*, and *Sec61b*) (Figure 8E).

### Metabolic and signaling pathways linked to macrophage activation m6A changes

Recent studies have highlighted a role for m6A in coordinating metabolic reprogramming and activation states in macrophages (Späth *et al*., 2025). To investigate if similar mechanisms operate during *L. amazonensis* infection, we examined a curated set of transcripts associated with immunometabolic pathways, including tricarboxylic acid (TCA) cycle, lipid metabolism, redox regulation, and inflammatory signaling (Supplementary Table 8).

Consistent with our global analyses, most transcripts retained at least part of their m6A landscape between conditions, indicating that infection does not induce widespread reorganization of m6A deposition. Instead, remodeling occurred selectively at specific transcripts and positions. Among the most dynamic examples, Socs3 and Map2k4 displayed marked increases in m6A site number, expanding from 3 to 11 and from 2 to 13 sites, respectively, highlighting extensive remodeling in transcripts involved in inflammatory feedback regulation and MAPK signaling. In contrast, several metabolic transcripts exhibited more moderate changes, including gains of m6A sites in *Hk1*, *Hk2*, *Ldlr*, and *Ogdh*, whereas *Sdha*, *Stat6*, *Slc25a1*, and *Tkt* showed site losses during infection (Supplementary Table 9).

To obtain a pathway-level view of these changes, we calculated a net remodeling score based on the balance between gained and lost m6A sites.

The strongest positive scores were observed for immune signaling pathways, followed by MAPK/stress signaling, lipid metabolism, glycolysis, and TCA-cycle-associated transcripts (Figure 9A). In contrast, redox-related pathways remained largely stable, while pentose phosphate pathway components exhibited a slight net loss of m6A sites.

**Figure 9.**
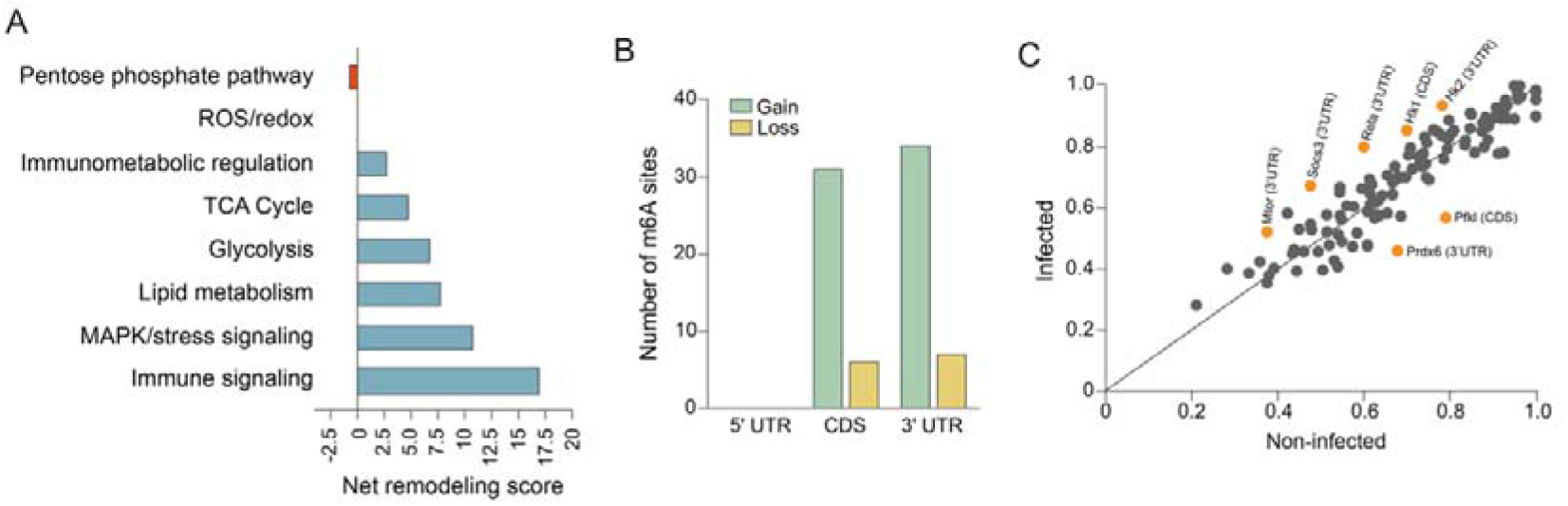
Selective m6A remodeling of immunometabolic transcripts during *L. amazonensis* infection. **A.** Net remodeling score across functional categories associated with macrophage activation and immunometabolic regulation. Scores were calculated as the difference between the number of gained and lost m6A sites within each pathway category, revealing preferential remodeling of transcripts involved in immune signaling, MAPK/stress signaling, lipid metabolism, glycolysis, and TCA cycle regulation. Negative values indicate a predominance of site loss. **B.** Distribution of gained and lost m6A sites across transcript regions. Newly acquired m6A sites were predominantly located within CDS and 3′ UTR, whereas no gain or loss events were detected in 5′ UTRs. **C.** Scatter plot comparing m6A mod ratio at sites shared between non-infected (x-axis) and infected (y-axis) macrophages. Each point represents an individual m6A site detected in both conditions. The diagonal line represents the line of identity, indicating equal m6A occupancy between conditions. Linear regression analysis revealed a strong correlation between conditions (R^2^ = 0.8481, *p* < 0.0001). Orange points highlight selected sites exhibiting the largest infection-associated changes in m6A occupancy, including sites within *Socs3*, *Rela*, *Mtor*, *Hk1*, *Hk*2, *Pfkl,* and *Prdx6* transcripts. Highlighted sites were selected based on the largest absolute differences in Δm6A mod ratio between infected and non-infected conditions.

Within the context of site gain and loss dynamics, newly acquired m6A sites were almost exclusively located within coding sequences and 3′ UTRs, whereas no gain or loss events were detected in 5′ UTRs (Figure 9B). Specifically, 31 gained sites were identified in CDS regions and 34 in 3′ UTRs, compared with only 6 and 7 lost sites, respectively.

Finally, we assessed changes in m6A occupancy at conserved sites. Overall, m6A levels remained highly correlated between non-infected and infected macrophages, indicating substantial preservation of site occupancy (Figure 9C). Nevertheless, several sites deviated from this global trend. Increased m6A occupancy was observed in transcripts including *Hk1*, *Hk2*, *Mtor*, Socs3, and Rela, whereas reduced occupancy was detected in *Pfkl* and *Prdx6*.

Although these changes were modest in magnitude, their concentration within key immunometabolic regulators suggests that m6A may contribute to fine-tuning macrophage responses during infection.

## Discussion

This study provides the first characterization of the host m6A epitranscriptome during *Leishmania* infection, revealing that *L. amazonensis* remodels the m6A regulatory machinery, elevates global RNA methylation, and drives transcriptome-wide shifts in modification distribution and site occupancy across thousands of host transcripts. The selectivity of this remodeling, preferentially targeting immune-regulatory and immunometabolic networks rather than globally restructuring the transcriptome, argues that the host epitranscriptome is not a passive bystander of infection but an active regulatory interface that is specifically engaged during the establishment of the intracellular *Leishmania* niche.

The consistent upregulation of METTL3 and downregulation of ALKBH5 across three independent patient transcriptomic datasets, validated in a dedicated cohort of 50 cutaneous leishmaniasis patients and recapitulated in RAW264.7 macrophages in vitro, establishes m6A machinery remodeling as a reproducible feature of leishmaniasis rather than a cell-line artifact. This METTL3-up/ALKBH5-down signature has been reported across phylogenetically distinct infection contexts, including septic cardiomyopathy and protozoan infection (Xia *et al*., 2021b; Wang *et al*., 2022a; Gong *et al*., 2023), suggesting it may represent a conserved host epitranscriptomic response to intracellular challenge. Mechanistically, the ALKBH5 downregulation in *C. parvum*-infected epithelial cells is driven by NF-κB p65 recruitment to the *Alkbh5* promoter downstream of TLR/MyD88 signaling (Xia *et al*., 2021b), a pathway that *Leishmania* also activates (Bichiou *et al*., 2021), making analogous NF-κB mediated eraser repression a plausible mechanistic hypothesis for future investigation.

The functional consequences of ALKBH5 loss during infection are context dependent. In *P. aeruginosa*-infected RAW264.7 macrophages, ALKBH5 knockdown amplifies expression of TLR4 pathway components by stabilizing their m6A-marked transcripts (Feng *et al*., 2022). In viral infection, ALKBH5 promotes nuclear retention of antiviral transcripts to sustain interferon responses (Gokhale *et al*., 2016). The reduction of ALKBH5 during *L. amazonensis* infection may therefore be detrimental to the parasite interest, allowing m6A marks to persist on immune-regulatory mRNAs, and could permit a partial, self-limiting inflammatory response that preserves the host cell while failing to clear the parasite. Also, our data reveal downregulation of METTL14, YTHDC1, and YTHDF3 in cutaneous leishmaniasis patients. The reduction in YTHDF3, which cooperates with both YTHDF1 and YTHDF2 and participates in functional redundancy during macrophage LPS responses (Balacco and Soller, 2019; Tong *et al*., 2021), adds a reader-level dimension to the machinery remodeling that extends beyond changes in modification abundance alone.

Elevated global m6A levels were detectable at 0 hpi (the earliest post-internalization time point) indicating that epitranscriptomic remodeling begins during the parasite-host contact phase itself, not as a delayed secondary response. The subsequent biphasic dynamics (peak at 4 hpi, partial normalization by 24 hpi) closely parallel the kinetics of m6A remodeling reported in *P. aeruginosa*-infected RAW264.7 macrophages (Feng *et al*., 2022), reinforcing the view that rapid epitranscriptomic engagement is a conserved feature of innate immune activation rather than a leishmaniasis-specific phenomenon. This early response is grounded in the convergence of rising METTL3 protein abundance and declining ALKBH5 levels between 2 and 8 hpi, creating a temporal window of unopposed m6A writing that directly accounts for the observed global methylation increase.

Transcriptome-wide m6A mapping by DRS demonstrated a 48% increase in modified transcripts and an 89% increase in high-confidence sites during infection, with 1,054 transcripts and 7,206 sites appearing exclusively in the infected state. Critically, this expansion is accompanied by increased modification occupancy at conserved sites (median m6A ratio 0.7429 vs. 0.7316; paired Wilcoxon, *p* = 0.0007), meaning that infection simultaneously broadens the methylated transcriptome and deepens methylation at positions already targeted in the basal state. The biological relevance of this increased occupancy is supported by Zhou et al. (2024), who showed that YTHDF2-mediated mRNA decay in the CDS is directly proportional to m6A site occupancy, where transcripts with higher modification levels are degraded more efficiently (Zhou *et al*., 2024). Taken together, these patterns point to a selective, infection-driven expansion of the m6A landscape rather than a non-specific global response, consistent with the targeted remodeling recently reported during *T. gondii* and *P. aeruginosa* infection (Feng *et al*., 2022; Qin *et al*., 2025b).

Infection nearly tripled the number of transcripts simultaneously modified in all three regions (47 to 140) and nearly doubled CDS–3′ UTR co-modification (743 to 1,306). This expansion of multi-region methylation is mechanistically significant because modifications in different regions recruit distinct reader proteins with opposing regulatory outputs. For example, 5′ UTR m6A promotes cap-independent translation under stress, CDS m6A drives YTHDF2-mediated decay (Zhou *et al*., 2024), and 3′ UTR m6A can either accelerate turnover via YTHDF2 or stabilize transcripts via IGF2BP-family readers depending on cellular context (Edupuganti *et al*., 2017; Shi *et al*., 2019). The co-occurrence of modifications across regions in the same transcript therefore substantially amplifies post-transcriptional regulatory complexity beyond what transcript abundance measurements capture (Chaparro *et al*., 2020). In line with this, infected samples also showed a slight increase in m6A density near the start codon. A similar, though more pronounced, enrichment near the start codon was reported in the spleen of *Plasmodium yoelii*-infected mice (Wang *et al*., 2022b), suggesting that start-codon-proximal methylation may be a recurring feature of the host response to protozoan infection.

Seven immune-related transcripts acquired m6A exclusively during infection: *Tlr4, Ccl2, Bcl3, Fosl2, Hmox1, Insig1*, and *Atp6v1g1*. The de novo methylation of *Tlr4* is particularly interesting given that METTL3-mediated m6A on *Tlr4* mRNA enhances its translation and amplifies TLR4/NF-DB signaling in macrophages (Liu *et al*., 2019), and that m6A occupancy at TLR4 pathway components correlates positively with gene expression during bacterial infection (Feng *et al*., 2022). Among shared immune transcripts, condition-specific sites acquired during infection showed significantly higher occupancy than those unique to controls (mean m6A mod ratio 0.5955 vs. 0.4829; *p* = 0.0206), and the most remodeled transcripts span activating factors (*Cd14, Irf5, Rela*) alongside negative regulators (*Socs3, Tgfb1, Tollip*). This distribution is consistent with the mixed M1/M2 macrophage phenotype and the balanced inflammatory state that sustains *Leishmania* persistence without triggering host cell death (Bichiou *et al*., 2021; Späth *et al*., 2025).

Within the immunometabolic transcriptome, the most extensive site-level remodeling occurred in MAPK and immune signaling components, with *Map2k4* acquiring 11 additional sites and *Socs3* expanding from 3 to 11. The *Map2k4* finding is directly connected to prior functional work demonstrating that YTHDF2-mediated decay of *Map2k4* and *Map4k4* mRNAs in LPS-stimulated RAW264.7 macrophages limits MAPK/NF-κB activation and constrains cytokine production (Yu *et al*., 2019). Increased m6A site density at *Map2k4* during *Leishmania* infection could modulate the efficiency of this YTHDF2-mediated decay, with downstream consequences for the JNK and p38 signaling that shape macrophage activation state. Elevated occupancy at shared sites in *Hk1, Hk2, Mtor*, and *Rela* further implicates m6A remodeling in the glycolytic and inflammatory reprogramming of infected macrophages, consistent with the IGF2BP2-dependent Warburg-like metabolic shift recently characterized in *L. amazonensis*-infected cells (Späth *et al*., 2025). Three newly acquired m6A sites in the 3′ UTR of *Igf2bp2* itself raise the possibility that m6A deposition on this transcript stabilizes IGF2BP2 mRNA, amplifying its availability to drive downstream glycolytic transcript stabilization, a mechanism that needs further investigation.

Perhaps the most unexpected finding was the de novo acquisition of multiple m6A sites on *Dicer1*, a transcript with no detectable methylation in non-infected macrophages. DICER1 sits at the center of miRNA biogenesis and translational repression, and its cleavage by the *L. donovani*-derived metalloprotease gp63 has been directly shown to impair host miRNA processing and increase parasite survival (Ghosh *et al*., 2013).

The acquisition of m6A on the *Dicer1* transcript during *L. amazonensis* infection suggests that epitranscriptomic regulation of this same host factor may represent a parallel or complementary mechanism of DICER1 control, potentially altering its translational efficiency or stability independently of proteolytic cleavage.

In conclusion, we show for the first time that *Leishmania* infection remodels the m6A landscape of host macrophages, and that this remodeling affects transcripts important for immune response, opening a new dimension of host-parasite biology that remained unexplored and maybe reveal new targets for host-directed therapeutic intervention.

## Materials and Methods

### Ethics statement

Human samples from patients diagnosed with cutaneous leishmaniasis were obtained through a previously approved study conducted at Instituto Aggeu Magalhães (IAM) - Fiocruz Pernambuco (CAAE: 11083812.7.0000.5190). Detailed clinical characteristics of the study population, diagnostic criteria, and sample collection procedures were previously described by Freitas e Silva et al., 2020 (de Freitas E Silva *et al*., 2020). All study procedures were approved by the appropriate Institutional Research Ethics Committee, and written informed consent was obtained from all participants prior to their enrollment.

### Public RNA-seq datasets analyses

Publicly available RNAseq datasets were retrieved from the Gene Expression Omnibus (GEO) and the Sequence Read Archive (SRA) to evaluate the expression profile of genes encoding m6A regulatory components (“writers”, “erasers” and “readers”). The selected datasets comprised transcriptomic data derived from skin lesion biopsies and whole blood samples from infected and non-infected individuals. A detailed description of all datasets included in the analysis is provided in Supplementary Table S1.

Raw sequencing data were processed using the Galaxy platform (https://usegalaxy.org) following a standardized RNA-seq analysis workflow. Gene expression levels were quantified and normalized using the Reads Per Kilobase per Million mapped reads (RPKM) and log₂-transformed for subsequent analyses. To ensure consistency across datasets, transcript lengths were retrieved using the RefSeq accession designated as canonical by UniProt for each gene, considering the most recent accession version and the complete mature transcript length, including the 5′ UTR, CDS, and 3′ UTR.

### Cell culture and macrophage differentiation

RAW264.7 murine macrophage-like cells (ATCC #TIB-71) were maintained in RPMI 1640 medium (Gibco, #31800022) supplemented with 10% heat-inactivated fetal bovine serum (FBS), 2 g/L sodium bicarbonate, 2.38 g/L HEPES, penicillin (100 U/mL), and streptomycin (133 μg/mL). Cells were maintained at 37 °C in a humidified atmosphere containing 5% CO₂.

THP-1 human monocytic cells (ATCC #TIB-202) were maintained in suspension in supplemented RPMI 1640 medium at 37 °C and 5% CO₂. Differentiation into macrophage-like cells was induced by treatment with 40 ng/mL phorbol-12-myristate-13-acetate (PMA) (Sigma-Aldrich, #P1585-1MG) for 72 h.

### Parasite culture

*L. amazonensis* promastigotes (MHOM/BR/1973/M2269) were maintained at 26 °C in M199 medium (Thermo Life Science, #31-100-019), pH 7.4, supplemented with 40 mM HEPES, 0.1 mM adenine, 4.62 mM NaHCO₃, 0.0001% biotin (Sigma, #B329), 10% heat-inactivated FBS, and penicillin/streptomycin.

### In vitro infection assays

RAW264.7 macrophages were seeded at 1 x 10□ cells per well in 24-well plates and infected with stationary-phase *L. amazonensis* promastigotes at a multiplicity of infection (MOI) of 10:1 parasites per cell. After 2 h of parasite– macrophage interaction, non-internalized parasites were removed by repeated PBS washes, and infected cultures were maintained under standard culture conditions. The 2 h macrophage parasite exposure was defined as 0 h post-infection (hpi). Samples were then harvested at 0, 2, 4, 8, and 24 hpi.

For THP-1 infections, differentiated macrophage-like cells were seeded at 1.5 x 10□ cells per culture flask and infected using the same MOI and experimental design. After 3 h of parasite exposure, non-internalized parasites were removed, and samples were collected at the indicated post-infection time points for downstream analysis.

### RT-qPCR analysis of m6A machinery components

Total RNA was extracted using TRIzol reagent (Thermo Fisher, #15596026) from peripheral blood mononuclear cells (PBMC) isolated from 50 patients diagnosed with cutaneous leishmaniasis and 10 healthy individuals. RNA concentration and purity were assessed using a NanoDrop ND-100 spectrophotometer (Thermo Fischer Scientific), and only samples presenting A260/280 and A260/230 absorbance ratios between 1.8 and 2.0 were used for downstream analyses.

cDNA synthesis was performed using Script 3.0 Reverse Transcriptase (Cellco, #PRT-102XS), according to the manufacturer’s instructions. RT-qPCR was performed using SYBR™ Green chemistry (Thermo Fisher Scientific, #4367659) in a ViiA™ 7 Real-Time PCR System (Thermo Fisher Scientific). Relative transcript abundance was determined using the comparative ΔCt method (Schmittgen and Livak, 2008), with *ACTIN* serving as the endogenous reference gene for normalization. Primer sequences used in these assays are listed in Supplementary Table S2.

### Western blot assays

Protein expression levels of METTL3 and ALKBH5 were evaluated in RAW264.7 macrophages collected at 0, 2, 4, 8, and 24 hpi. Total protein extracts were prepared using RIPA lysis buffer supplemented with a protease inhibitor cocktail, and protein concentration was determined using the BCA Protein Assay Kit (Thermo Scientific, #23227), according to the manufacturer’s instructions. Equal amounts of proteins (30 µg per sample) were resolved by 10% SDS-PAGE and transferred onto nitrocellulose membranes. Membranes were blocked in an appropriated blocking buffer and incubated overnight at 4 °C with primary antibodies against METTL3 (rabbit, 1:1,000; Abcam, #Ab195352) or ALKBH5 (rabbit, 1:1,000; Novus Biologicals, #NBP1-82188), followed by incubation with HRP-conjugated secondary antibodies (Sigma-Aldrich, #A9044). GAPDH (BioLegend, #649202) was used as an internal loading control. Protein signals were detected using the Li-Cor Odyssey imaging system, and relative protein abundance was quantified by densitometric analysis.

### Immunofluorescence analysis

Immunofluorescence assays were performed to evaluate the abundance and subcellular distribution of METTL3 and ALKBH5 in infected RAW264.7 macrophages. Cells were fixed with 4% paraformaldehyde in PBS, permeabilized with 0.5% Triton X-100, and blocked with PGN–saponin solution (0.25% porcine skin gelatin and 0.10% saponin in PBS) to minimize nonspecific antibody binding. Samples were incubated with primary antibodies against METTL3 or ALKBH5 followed by species-specific Alexa Fluor–conjugated secondary antibodies (anti-mouse IgG Alexa Fluor™ Plus 488 for METTL3 and anti-rabbit IgG Alexa Fluor™ Plus 647 for ALKBH5). Cell nuclei were stained with Hoechst 33342 (Thermo Fisher Scientific, ##34580). Images were acquired using a Leica TCS SP5 II confocal microscope. Fluorescence intensity was quantified at the whole-cell level using ImageJ software with a minimum of 100 cells analyzed per condition.

### Dot blot analysis for global m6A quantification

Global m6A levels were assessed in total RNA extracted from RAW264.7 macrophages collected at 0, 2, 4, 8, 24 hpi. Equal amounts of RNA were subjected to dot blot assays using a specific anti-m6A antibody (Abcam, #ab151230), as previously described in (Chen *et al*., 2021). Briefly, RNA samples were denatured, spotted onto nylon membranes, and immobilized prior to immunodetection. Membranes were then incubated with the anti-m6A antibody, followed by appropriate secondary antibodies for signal detection. Chemiluminescent signals were detected using the Li-Cor Odyssey imaging system, and relative m6A abundance was determined by densitometric analysis considering the methylene blue signal as loading control.

### Direct RNA sequencing (DRS)

For Direct RNA Sequencing analyses, RAW264.7 macrophages were infected with stationary-phase *L. amazonensis* promastigotes, and samples were collected at 8 hpi for RNA extraction. Total RNA was isolated using TRIzol reagent followed by purification with the PureLink RNA Mini Kit (Invitrogen, #12183018A). RNA purity and concentration were verified via NanoDrop spectrophotometry and agarose gel electrophoresis prior to library preparation.

DRS libraries were prepared using 100 ng of total RNA derived from non-infected and infected macrophages using the SQK-RNA004 kit (Oxford Nanopore Technologies, ONT, Oxford, UK). Libraries were loaded onto R10.4.1 flow cells (FLO-MIN106) after confirmation of a minimum of 800 active nanopores per flow cell. Sequencing was performed on a MinION1 Mk1B device using MinKNOW software (v25.09.16), under standard Oxford Nanopore acquisition settings. Individual sequencing runs were performed for 24-48 h to maximize data yield and transcriptome coverage.

### DRS data analysis and m6A identification

Raw pod5 files were basecalled using Dorado (https://github.com/nanoporetech/dorado), through the high-accuracy (hac) mode, retaining only reads with an average Phred quality score ≥ 10.

Sequencing quality control was assessed using the manufactureŕs MinKNOW® platform (v25.09.16). Across all libraries, an average of 491 thousand reads were obtained per sample, with 92.6% mean of uniquely mapped reads against the reference transcriptome (see below). FASTQ files generated from individual libraries were concatenated using catfishq (https://github.com/philres/catfishq) and mapped against the *Mus musculus* reference transcriptome (GRCm39, Ensembl release 111) using the minimap2 aligner (v2.28) (Li, 2018), with thefollowing parameters “-ax map-ont -uf -k14 --secondary=no”. The sorted BAM files generated were used for transcript quantification using NanoCount (Gleeson *et al*., 2022) with the --extra_tx_info option enabled.

For m6A identification, nanopore raw signal data were processed using f5c v1.6 index (--slow5 ${sample}.blow5 ${sample}.fastq.gz) and eventalign (--kmer-model rna004.5mer.model --pore rna004 --slow5 ${sample}.blow5 --scale-events --signal-index –rna -r ${sample}.fastq.gz -b ${sample}.bam -g Mus_musculus.GRCm39.cdna.all.fa, followed by m6anet (v2.1) analysis (modules dataprep and inference --num_iterations 1000) (Hendra *et al*., 2022). m6A sites were identified based on a modification probability (or confidence) threshold ≥ 0.80.

To increase robustness and reproducibility, m6A-calls from three biological replicates of each experimental condition (non-infected and infected macrophages) were integrated prior to downstream analyses. Redundant entries were removed based on transcript identifier (Transcript_ID) and m6A-site position. Two complementary analytical approaches were subsequently applied: 1) transcript-centric analyses collapsed all modification events within a transcript to determine the total number of unique m6A-modified transcripts; and 2) site-centric analyses retained transcript position information to quantify the total number of individual m6A sites detected in each condition.

Transcript annotation was performed using the Ensembl BioMart platform to retrieve gene identifiers (gene ID), transcript length, transcript biotype, and isoform descriptions. m6A site density (number of sites per transcript) was calculated as the number of m6A sites detected by transcript. The m6A modification ratio (m6A mod_ratio), ranging from 0 to 1, was used as a proxy for site occupancy and corresponds to the fraction of methylated reads relative to the read coverage at a given position.

To investigate infection-associated epitranscriptomic remodeling within immune-related pathways relevant to *Leishmania* infection, a manually curated reference gene set was generated based on an extensive literature review of host immune responses to *Leishmania* infection. Relevant studies were identified through systematic searches of the PubMed using the terms “*Leishmania*”, “leishmaniasis”, and “immune response”. This approach resulted in a curated panel of 333 genes implicated in macrophage activation, innate immune signaling, inflammatory responses, antigen presentation, cytokine regulation, and host defense mechanisms during leishmaniasis. Corresponding versioned transcript identifiers were retrieved using the Ensembl BioMart and integrated with the DRS-derived m6A dataset for downstream analyses.

Functional enrichment analyses were performed using the STRING database (version 12.0) (Szklarczyk *et al*., 2025), including KEGG pathway annotation and Gene Ontology analyses. Statistical significance was determined using the default enrichment parameters implemented in STRING, with terms considered significant at an FDR-adjusted p-value < 0.05.

## Statistical analysis

All statistical analyses were performed using GraphPad Prism 8.0. Data distribution was assessed prior to statistical testing to determine the appropriate analytical approach. Comparisons between two independent groups were conducted using an unpaired Student’s *t*-test for normally distributed data (parametric) or the Mann–Whitney *U* test for non-normally distributed data (non-parametric). Paired datasets were analyzed using the Wilcoxon matched-pairs signed-rank test. Differences in cumulative data distributions were evaluated using the Kolmogorov–Smirnov test. For comparisons involving more than two groups, statistical significance was determined using the Kruskal–Wallis test followed by Dunn’s post-hoc test when applicable. Unless otherwise stated, a two-tailed p-value < 0.05 was considered statistically significant. The specific statistical test applied, sample sizes (n), and definitions of biological and technical replicates are indicated in the corresponding figure legends.

## Acknowledgements

This study was financed, in part, by the São Paulo Research Foundation (FAPESP), Brasil. Process Number 2022/03075-0 to N.M.; 2025/06837-7 to AGM; 2025/06325-6 and 2023/02341-1 to AHR; 2025/03898-5 to MAGN; to 2025/06986-2 to FCM; NM and RLAMN are CNPq research fellow (grant numbers CNPq 306191/2024-5 and 312353/2023-5, respectively). Research coordinated by RLMN is supported by grants from Fapemig (grant number #RED-00196-23); and “Programa Inova Fiocruz” from Oswaldo Cruz Foundation. Authors thank the Genomics Platform, RPT01E; Real-time and digital PCR platform, RPT09D; part of the Technological Platforms Network, Oswaldo Cruz Foundation, Fiocruz.Computing-intense data analyses from this work were undertaken on the AIRE cluster, part of High-Performance Computing facilities at the University of Leeds, UK, who we thank for regular maintenance and technical support. We also thank Marcelo Briones for assistance with Nanopore sequencing and Clara Lúcia Barbiéri Mestriner, Patrícia Xander Batista, Rogéria Cristina Zauli, João Paulo Ferreira Rodrigues and Leonardo Loch for technical assistance. The authors thank the members of the Laboratório de Biologia Molecular de Patógenos (LBMP) for valuable discussions.

## Author contributions

Conceptualization, A.G.M. and N.S.M.; Methodology, A.G.M., M.A.N.G., C.E.A., B.S.B., A.H.R., P.O.L.M., and N.S.M.; Software, M.A.N.G., C.E.A., and E.J.R.V.; Validation, P.O.L.M. and E.J.R.V; Formal analysis, A.G.M., M.A.N.G., C.E.A., and E.J.R.V.; Investigation, A.G.M.; P.O.L.M; B.S.B; and E.J.R.V.; Resources, P.O.L.M., R.F.S., E.J.R.V., and N.S.M.; Data curation, A.G.M. and E.J.R.V.; Writing – original draft, A.G.M. and N.S.M.; Writing – review & editing, A.G.M., R.F.S., E.J.R.V., and N.S.M.; Visualization, A.G.M. and N.S.M.; Supervision, N.S.M.; Project administration, N.S.M.; Funding acquisition, N.S.M.

## Data Sharing and Data Availability

All DRS raw data will be available prior publication at GEO or SRA databases.

## Disclaimer

The authors used AI tools to assist in language editing to improve the quality of reading.

