## Supplementary Figures for "Direct RNA sequencing reveals selective remodeling of the host m6A epitranscriptome during *Leishmania* infection"

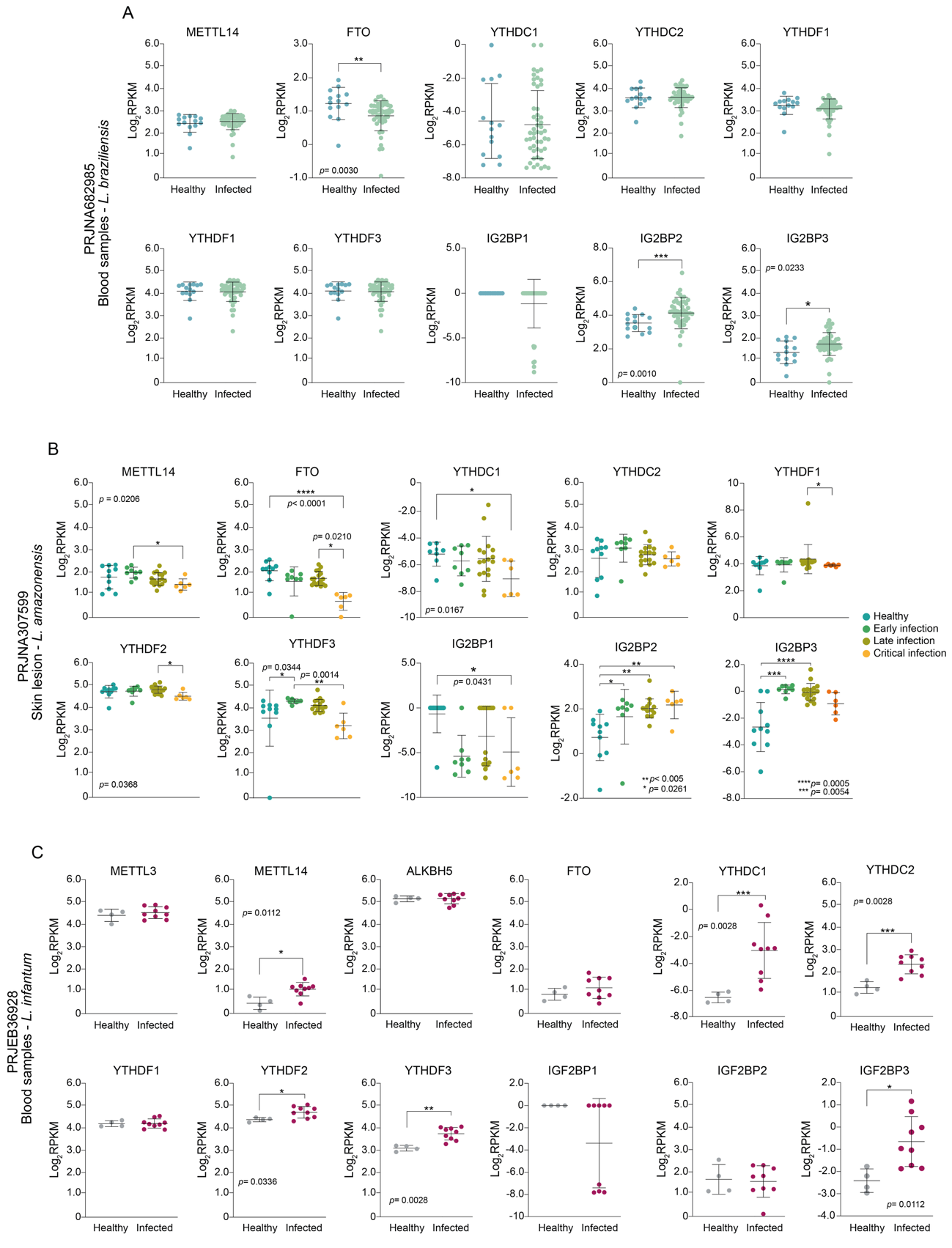

**Supplementary Figure 1. Expression profiles of additional m6A regulatory factors in publicly available leishmaniasis transcriptomic datasets. A.** Expression of m6A regulatory components in whole-blood samples from healthy individuals and patients with cutaneous leishmaniasis caused by *L. braziliensis* (GEO: GSE162760). **B.** Expression of m6A regulatory components in skin lesions biopsies from patients infected with *L. amazonensis* stratified according to disease stage (early infection, late infection, and chronic infection from PRJNA307599). **C.** Expression of m6A regulatory components in whole-bloody samples from healthy individuals and patients with visceral leishmaniasis caused by *Leishmania infantum* (PRJEB36928). The analyzed factors include the writer METTL14, the eraser FTO, the reader proteins YTHDC1, YTHDC2, YTHDF1, YTHDF2, and YTHDF3, as well as members of the IGF2BP family (IGF2BP1–3). Expression values were obtained from publicly available RNA-seq datasets and are presented as log₂-transformed RPKM values. Data are shown as mean ± SD. Statistical significance was determined using the tests implemented in the original analysis pipeline and is indicated in each panel (**p* < 0.05, ***p* < 0.01, ****p* < 0.001, *****p* < 0.0001).

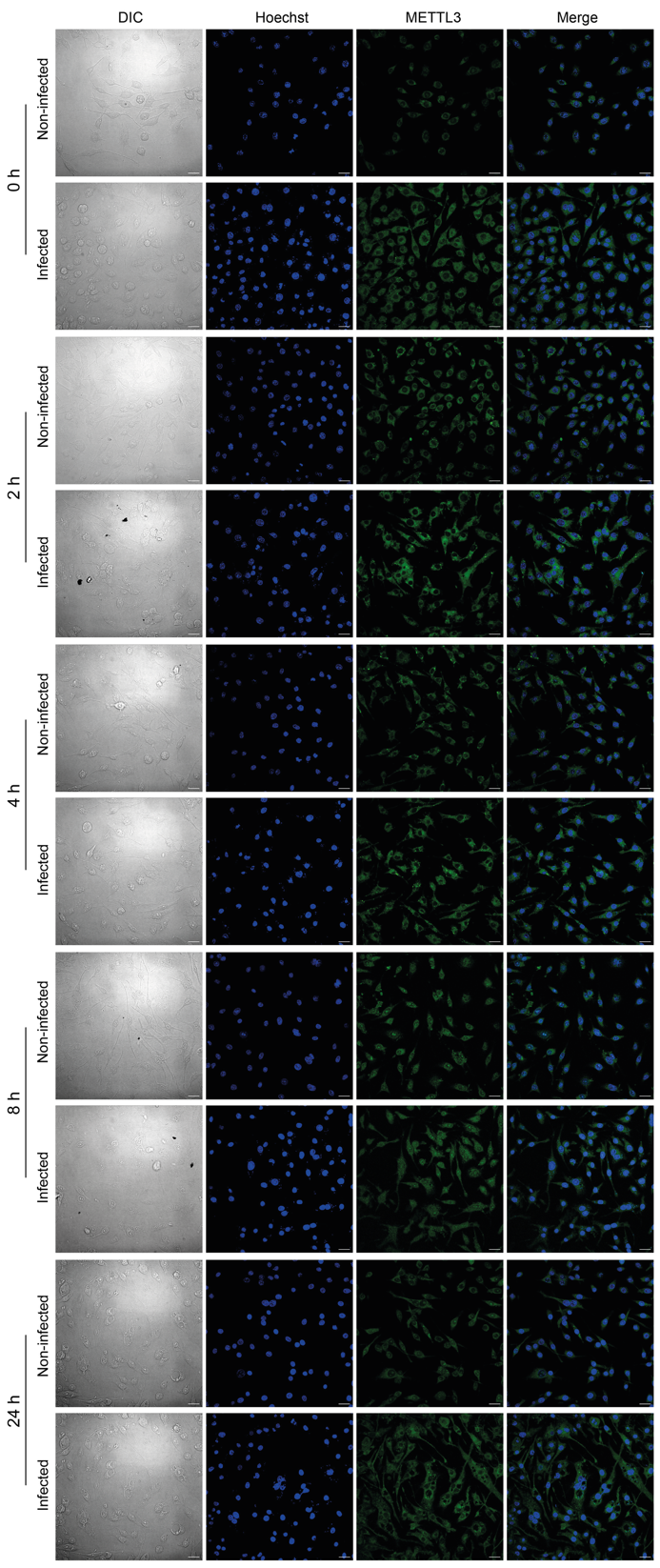

**Supplementary Figure 2. Immunofluorescence analysis of METTL3 during *L. amazonensis* infection.** Representative confocal microscopy images of RAW264.7 macrophages infected in vitro with *L. amazonensis* at 0, 2, 4, 8, and 24 h post-infection. METTL3 was detected by immunofluorescence (green), and nuclei were counterstained with Hoechst 33342 (blue). Scale bar, 19 µm.

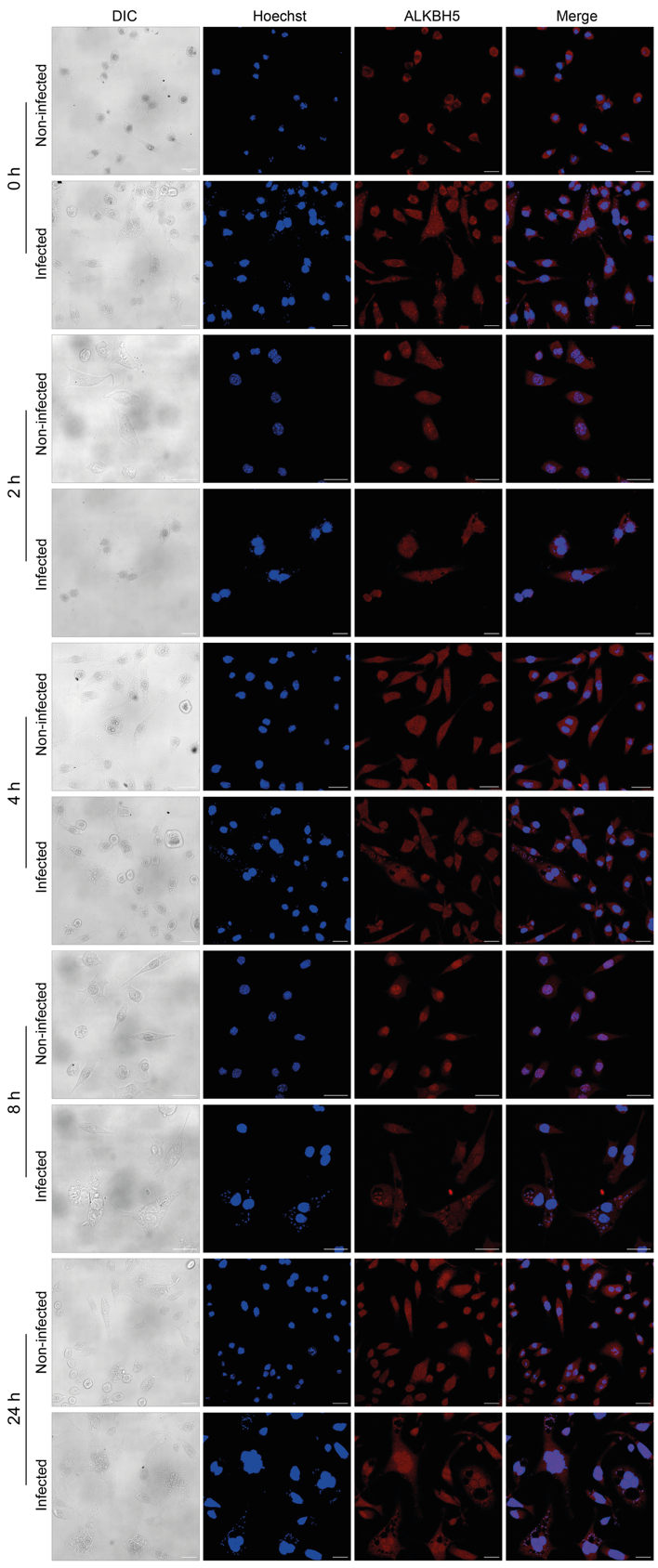

**Supplementary Figure 3. Immunofluorescence analysis of ALKBH5 during *L. amazonensis* infection.** Representative confocal microscopy images of RAW264.7 macrophages infected in vitro with *L. amazonensis* at 0, 2, 4, 8, and 24 h post-infection. ALKBH5 was detected by immunofluorescence (red), and nuclei were counterstained with Hoechst 33342 (blue). Scale bar, 19 µm.

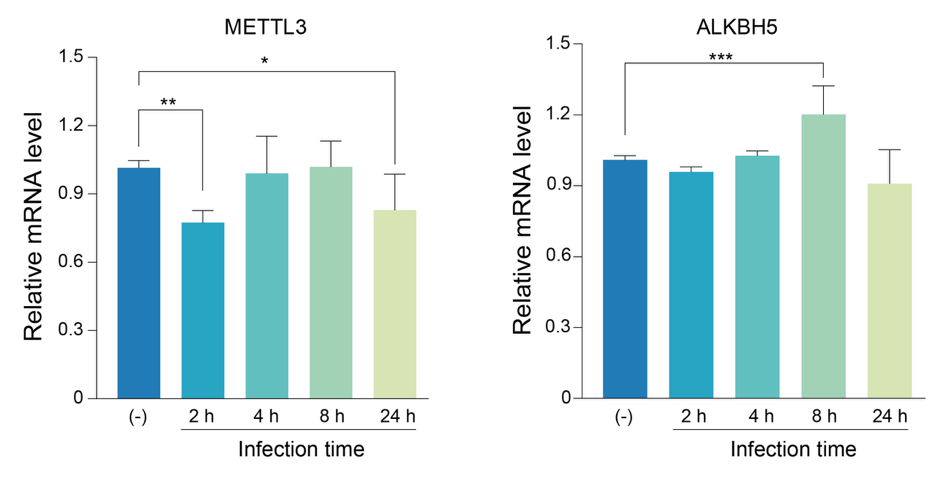

**Supplementary Figure 4. Modulation of METTL3 and ALKBH5 transcripts during *L. amazonensis* infection in macrophages.** Relative mRNA levels of METTL3 and ALKBH5 were quantified by RT-qPCR in macrophages with *L. amazonensis* promastigotes at 2, 4, 8 and 24 h post-infection. Gene expression levels were normalized to the endogenous *ACTIN* gene and expressed related to the non-infected condition (-). Data are presented as mean ± SD. Statistical significance was determined using one-way ANOVA with multiple-comparison testing (**p* < 0.05*, **p* < 0.01*, ***p* < 0.001*).*

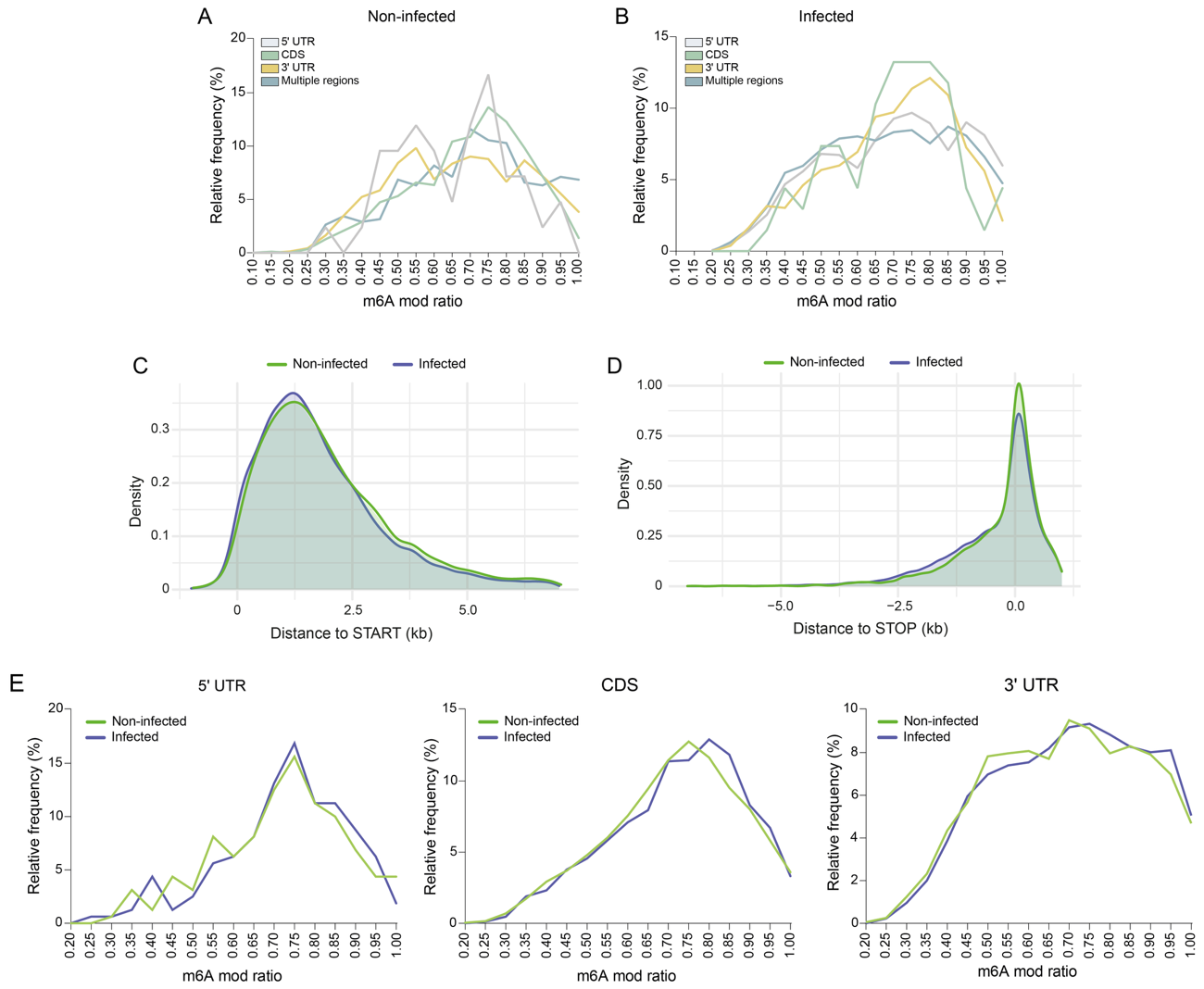

**Supplementary Figure 5. Regional and positional distribution of m6A modifications in non-infected and *L. amazonensis*-infected macrophages. A–B.** Frequency distributions of m6A mod ratio values for m6A sites located within the 5′ UTR, CDS, 3′ UTR, or in transcripts containing m6A sites across all three transcript regions (multiple regions), shown for non-infected (**A**) and infected (**B**) macrophages. **C.** Kernel density distribution of m6A sites relative to the translation start codon. **D.** Kernel density distribution of m6A sites relative to the translation stop codon. **E.** Frequency distributions of m6A mod ratio for m6A sites shared between non-infected and infected macrophages within the 5′ UTR, CDS, and 3′ UTR regions. Curves represent the relative frequency of shared sites across m6A mod ratio intervals, highlighting region-specific occupancy patterns between conditions.

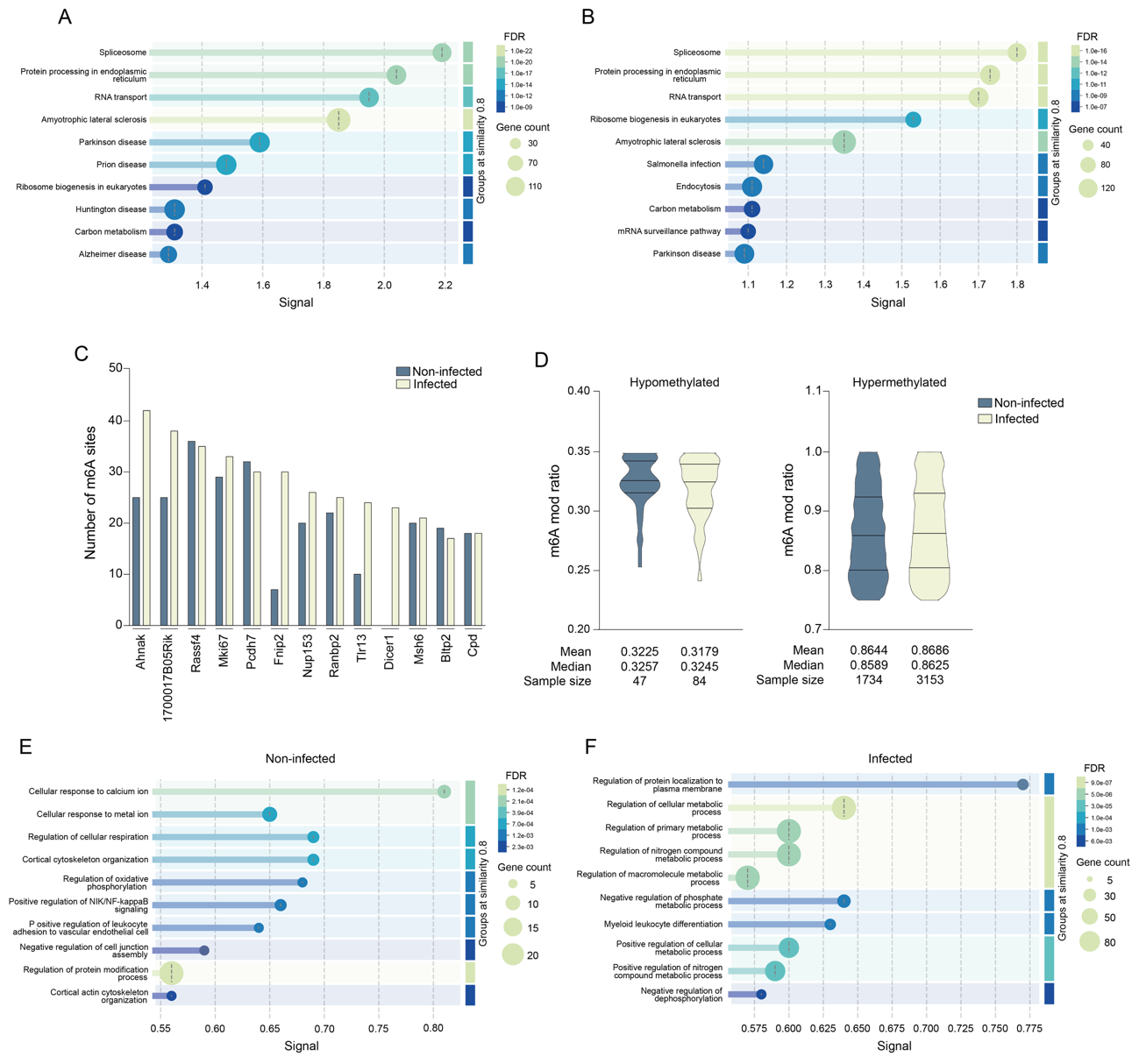

**Supplementary Figure 6. Functional characterization of m6A-modified transcripts identified in non-infected and *L. amazonensis*-infected macrophages. A.** KEGG pathway enrichment analysis of genes corresponding to m6A-modified transcripts identified in non-infected macrophages. **B.** KEGG pathway enrichment analysis of genes corresponding to m6A-modified transcripts identified in *L. amazonensis*-infected macrophages. Bubble size represents the number of genes associated with each pathway, and color indicates the false discovery rate (FDR). **C.** Top 13 transcripts containing the highest numbers of detected m6A sites in non-infected and infected macrophages. Bars indicate the total number of m6A sites identified in each transcript. **D.** Distribution of m6A modification ratios (m6A mod ratio) for stoichiometrically hypomethylated (m6A mod ratio ≤ 0.35) and hypermethylated (m6A mod ratio ≥ 0.75) sites in non-infected and infected macrophages. Violin plots display the distribution of m6A occupancy values, and summary statistics (mean, median, and sample size) are shown below each plot.  **E.** Gene Ontology – Biological Process enrichment analysis of genes corresponding to transcripts modified by m6A simultaneously in the 5′ UTR, CDS and 3′ UTR regions under the non-infected condition. **F.** Gene Ontology – Biological Process enrichment analysis of genes corresponding to transcripts modified by m6A simultaneously in the 5′ UTR, CDS and 3′ UTR regions under the infected condition.
